# Single administration of mosaic-8b RBD-nanoparticle vaccine prepared with atomic layer deposition technology elicits broadly neutralizing anti-sarbecovirus responses

**DOI:** 10.1101/2025.06.16.660007

**Authors:** Alexander A. Cohen, Jennifer R. Keeffe, Annie V. Rorick, Semi Rho, Ange-Celia Priso Fils, Lusineh Manasyan, Han Gao, Priyanthi N.P. Gnanapragasam, Hans H. Funke, Theodore W. Randolph, Robert L. Garcea, Pamela J. Bjorkman

## Abstract

Atomic layer deposition (ALD), a new vaccine technology, permits multiple dosing with a single administration by pulsatile release of one or more immunogens. We evaluated ALD delivery of mosaic-8b [60-mer nanoparticles presenting 8 different SARS-like betacoronavirus (sarbecovirus) receptor-binding domains (RBDs)] that elicits broadly cross-reactive antibodies and protects against mismatched sarbecoviruses not represented by RBDs on mosaic-8b. Compared with conventional prime-boost immunizations, ALD-delivered mosaic-8b RBD-nanoparticles elicited antibodies in both naïve and pre-vaccinated mice with improved mismatched binding and neutralization. Results of RBD epitope mapping of serum antibodies from ALD-delivered mosaic-8b were consistent with broader coverage of RBD epitopes compared to conventional immunizations, and systems serology revealed distinct IgG subclass and FcγR-binding IgG distributions. These results suggest that ALD is a promising technology for use with mosaic-8b RBD-nanoparticle vaccines to protect against future sarbecovirus spillovers and support applications for ALD vaccine delivery to elicit cross-reactive antibodies against rapidly mutating or diverse pathogens.

## INTRODUCTION

Vaccine technologies have recently advanced, as evidenced by the introductions of lipid-nanoparticles (LNP) as delivery vehicles for mRNA antigens and concurrent delivery of multiple antigens on single recombinant protein platforms.^1–3^ Despite the promise of these innovations, such formulations have barriers to widespread availability with respect to their storage conditions and the need for separate boosting doses for many antigens. We recently described a new protein vaccine technology that permits multiple prime-boost dosing with a single administration.^4–8^ This technology utilizes a highly scalable molecular deposition process (atomic layer deposition; ALD^9^) to coat the surface of thermostable microparticles containing vaccine immunogens with precise numbers of molecular layers of alumina that not only serve as adjuvant, but can be modulated to release the antigen at pre-determined time points after immunization. This technology enables temporally separated primer and booster vaccine doses from a single administration when immunogens coated with different numbers of alumina layers (and therefore released at different time points) are mixed into a single formulation^4^ (Figure 1A). The sequential release of the vaccine immunogen from a single injection is hypothesized to guide the developing B cell response toward progressive development of high-affinity specific memory B cells rather than by boosting with subsequent injections. In addition, higher neutralizing antibody (Ab) titers with the coated preparations have been observed than can be achieved with multiple immunization regimens of conventional formulations, a result which we hypothesize is the result of Ab maturation reflecting a higher affinity for the antigen at later times.^4,10^

**Figure 1.**
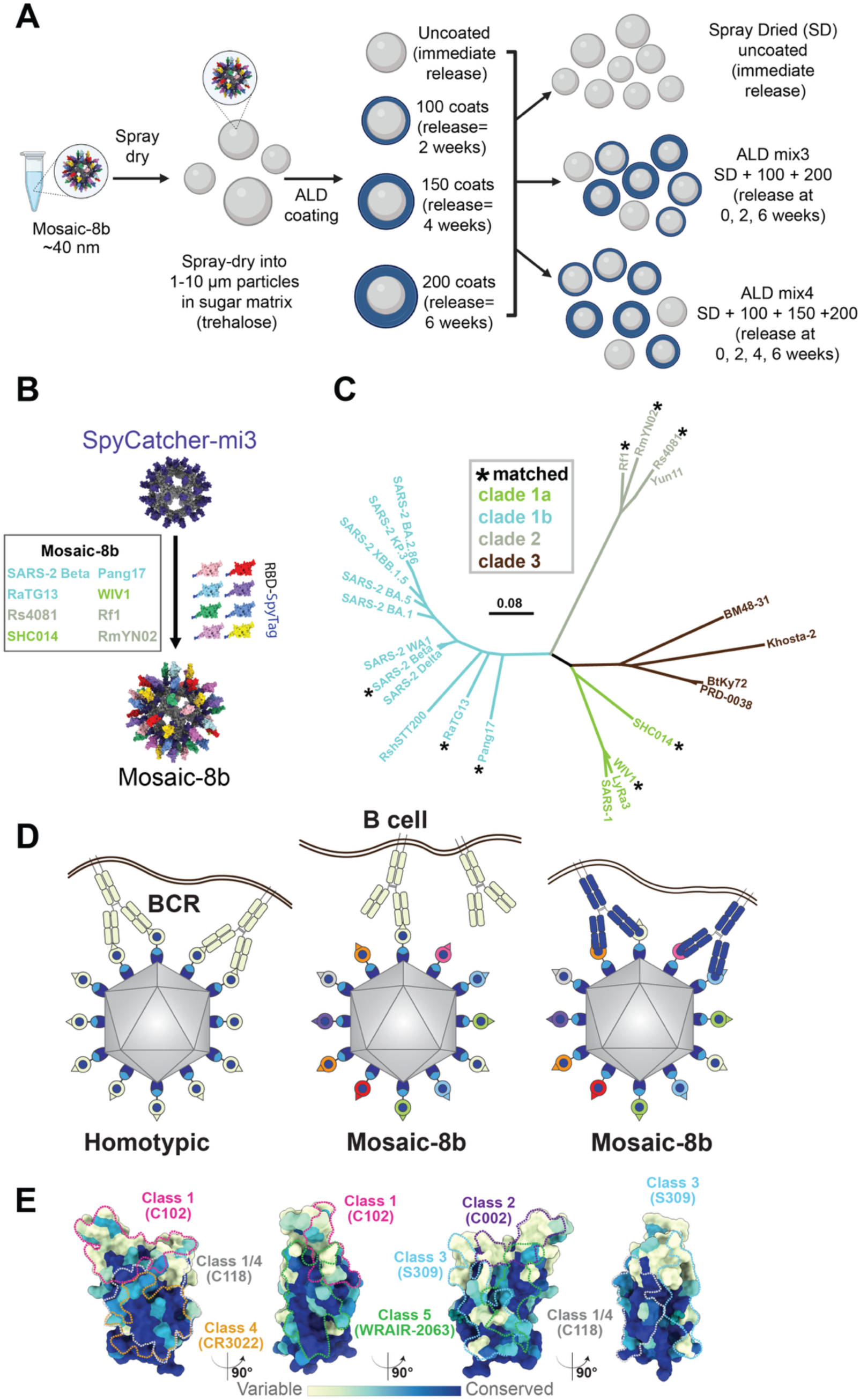
ALD formulation of mosaic-8b RBD-nanoparticles. (A) Preparation of the antigen using ALD. Mosaic-8b RBD-nanoparticles were linked to Spy-Catcher mi3^19^ and spray-dried in a buffer containing trehalose to form spherical ∼10 nm diameter particles. These spherical trehalose-antigen particles were then coated with precisely defined layer numbers of alumina using ALD. Resulting particles were then mixed according to the desired antigen release intervals.^4^ (B) Model of a mosaic-8b RBD-nanoparticle constructed using coordinates of an RBD (PDB 7BZ5), SpyCatcher (PDB 4MLI), and i3-01 nanoparticle (PDB 7B3Y). (C) Phylogenetic tree of selected sarbecoviruses calculated based on amino acid sequences of RBDs aligned using Clustal Omega.^43^ Tree was constructed using a Jukes-Cantor generic distance model with Geneious Prime® 2023.1.2. Viruses with RBDs included in mosaic-8b RBD-nanoparticles are indicated with an asterisk. The scale bar represents a phylogenetic distance of 0.08 nucleotide substitutions per site. (D) Avidity hypothesis schematic. Left: Membrane-bound BCRs use avidity to bind a strain-specific epitope (pale green triangle) on pale green antigens attached to a homotypic nanoparticle. Middle: BCRs cannot use avidity to bind a strain-specific epitope (triangle) on pale green antigen attached to a mosaic nanoparticle. Right: BCRs can use avidity to bind to common epitope (blue circle) presented on different antigens attached to a mosaic nanoparticle, but not to strain-specific epitopes (triangles). (E) Relative conservation of RBD epitopes. ConSurf^44^ calculation of sequence conservation of 16 sarbecovirus RBDs plotted on a surface representation of SARS-2 RBD (PDB 7BZ5). Anti-RBD class 1, 2, 3, 4, 1/4, and 5 Ab epitopes^22–25^ are outlined in different colored dots using epitope information from structures of representative Abs bound to SARS-2 spike or RBD (C102: PDB 7K8M; C002: PDB 7K8T, S309: PDB 7JX3; CR3022: PDB 7LOP; C118: PDB 7RKV; WRAIR-2063: PDB 8EOO).

The COVID-19 pandemic caused by the SARS-CoV-2 virus (hereafter SARS-2) prompted the development of many new vaccines,^11^ some or all of which potentially could be improved using ALD technology to broaden protection. We previously described a vaccine designed to protect against new SARS-2 variants and zoonotic SARS-like betacoronaviruses (sarbecoviruses) with spillover potential^12–15^ that could be delivered as a single-dose ALD immunization. The vaccine candidate, mosaic-8b, involves simultaneous display of eight different sarbecovirus receptor-binding domains (RBDs) arranged randomly on 60-mer nanoparticles^15^ (Figure 1B,C).

We hypothesized that mosaic RBD-nanoparticles would preferentially elicit cross-reactive Abs if B cell receptors (BCRs) on cross-reactive Ab-producing B cells are able to crosslink using both of their antigen-binding Fab arms between adjacent non-identical RBDs to recognize conserved epitopes, as compared with B cells presenting BCRs that bind to variable epitopes, which should only rarely crosslink between adjacent non-identical RBDs^14^ (Figure 1D; avidity hypothesis schematic). In contrast, homotypic RBD-nanoparticles presenting identical RBDs should predominantly bind BCRs that recognize immunodominant strain-specific epitopes (Figure 1D). We obtained evidence for this hypothesis using deep mutational scanning (DMS)^16^ to map epitopes recognized by IgGs in polyclonal antisera elicited by mosaic-8b versus homotypic SARS-2 RBD-nanoparticles: Abs from mosaic-8b antisera primarily targeted more conserved RBD epitopes, whereas Abs from homotypic antisera primarily targeted variable, and more accessible, epitopes^14^ (Figure 1E). When we evaluated responses against matched and mismatched viruses (i.e., represented or not represented by an RBD on the nanoparticle), we showed in animal models that mosaic-8b RBD-nanoparticles showed enhanced heterologous binding, neutralization, and protection from sarbecovirus challenges compared with homotypic (SARS-2 RBD only) nanoparticles.^14,15^ Broader Ab responses elicited by mosaic-8b compared with homotypic RBD-nanoparticles were found in both naïve animals and in animals that had been pre-vaccinated with COVID-19 vaccines.^13^

Here, we investigated whether improvements in vaccine-induced Ab responses using ALD pulsatile technology^4–6^ also could be applied to delivery of mosaic-8b RBD-nanoparticles. When compared to conventional mosaic-8b immunization (i.e., separate prime and boost injections), we found that a single administration of ALD vaccine mosaic-8b preparations with different intervals of antigen release (ALD mix3 or ALD mix4) elicited significantly higher neutralization titers than conventional mosaic-8b immunization against difficult-to-neutralize mismatched viral strains in both originally naïve and in pre-vaccinated mice. In addition, ALD-elicited Abs showed differences in epitope recognition and in their relative proportions of IgG subclasses compared with non-ALD conventional prime and boost injections. Together, our results demonstrate the ability of ALD coating of mosaic-8b RBD-nanoparticles, and likely other broad-based vaccines, to improve the quality of elicited Abs by inducing broader neutralizing responses.

## RESULTS

### Production of ALD and non-ALD formulated versions of mosaic-8b RBD-nanoparticles

To generate mosaic-8b RBD-nanoparticles, we used the SpyCatcher-SpyTag system^17,18^ to covalently attach RBDs (SARS-2 Beta RBD and seven other sarbecovirus RBDs) with C-terminal SpyTag sequences to a SpyCatcher-mi3 60-mer protein nanoparticle^19^ (Figure 1B,C). Mosaic-8b RBD-nanoparticles were then spray-dried in a buffer containing trehalose, followed by coating the resulting spherical microparticles with precisely defined numbers of alumina layers using ALD^4^ (Figure 1A).

ALD mix3 powder contained three formulations of antigen: mosaic-8b that was (i) spray-dried only (SD), (ii) spray-dried with 100 coats of alumina, and (iii) spray-dried with 200 coats, resulting in antigens released at ≈ 0, 2, and 4 weeks. ALD mix4 powder contained four formulations of antigen: mosaic-8b that was (i) spray-dried only, (ii) spray-dried with 100 coats of alumina, (iii) spray-dried with 150 coats of alumina, and (iv) spray-dried with 200 coats, resulting in release at ≈ 0, 2, 3, and 4 weeks (Figure 1A). When reconstituted, ALD mix3 and ALD mix4 each represented a single 5 µg RBD dose per animal, which is comparable to the total RBD content in the conventional mosaic-8b RBD-nanoparticle prime and boost injections (two 2.5 µg injections).

### Single ALD-formulated mosaic-8b immunizations elicited superior Ab responses compared with conventional prime-boost immunization

We first addressed whether pulsatile release of mosaic-8b using ALD-coated antigens improved the immunogenicity of mosaic-8b in naïve mice. We immunized mice with either a single dose of ALD mix3, ALD mix4, or spray dried mosaic-8b without ALD coating (SD) and compared these responses to responses elicited by a conventional immunization regimen (two doses of Addavax-adjuvanted mosaic-8b delivered 4 weeks apart) (Figure 2A). Serum samples at 12- and 31-weeks post-prime were analyzed by ELISA (Figure 2B) and pseudovirus neutralization assays (Figure 2C) to determine serum Ab binding and neutralization titers, respectively. The Ab ELISA binding data and neutralization results are presented in the form of line plots (ELISA) or box and whisker plots (neutralization) with determinations of statistically significant differences between cohorts evaluated using pairwise comparisons for each viral strain.

**Figure 2.**
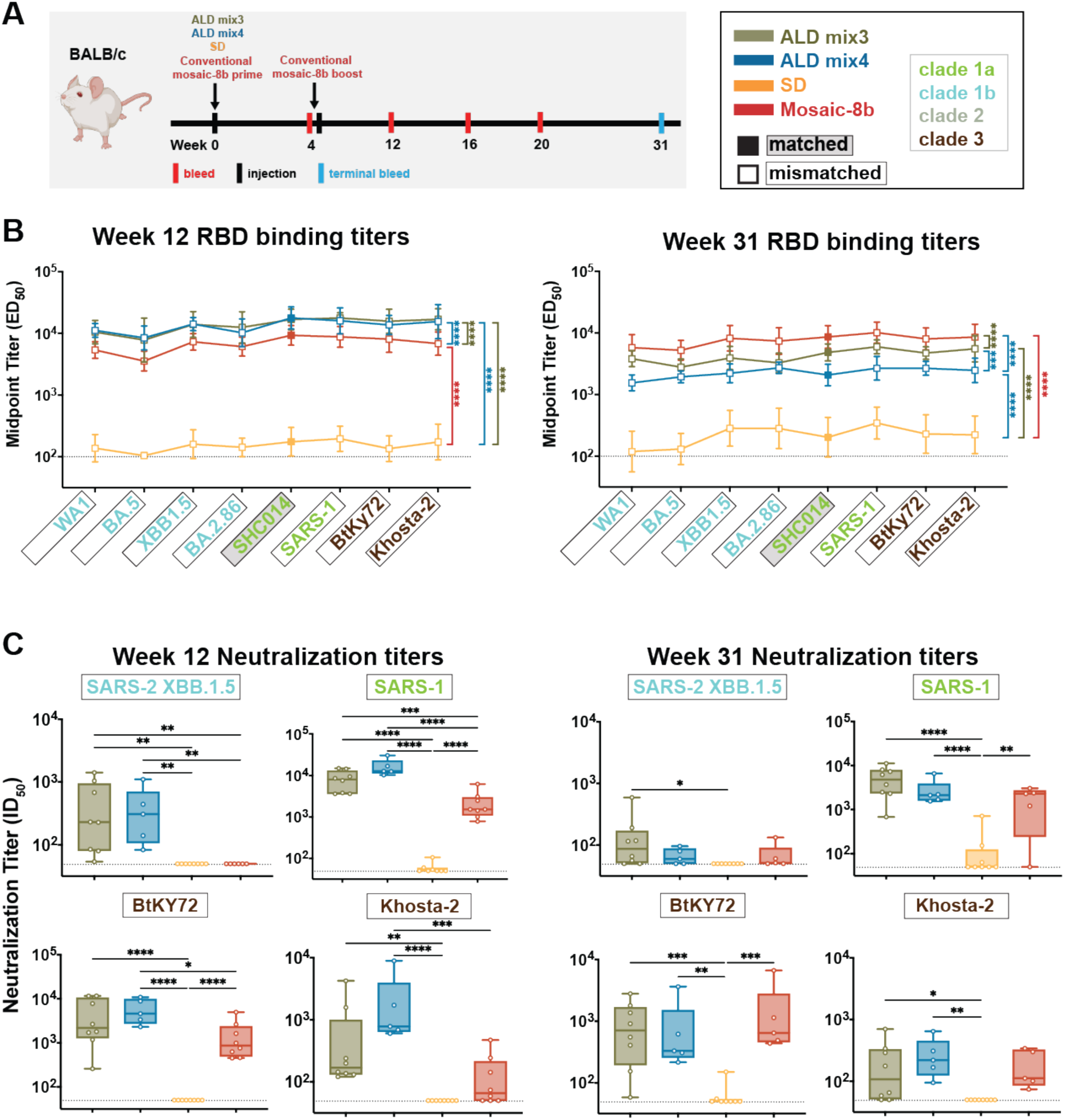
ALD-coated mosaic-8b immunization elicits more broadly neutralizing Ab responses than conventional mosaic-8b immunizations in originally naïve mice. Significant differences between cohorts linked by vertical lines in panels B and C are indicated by asterisks: p<0.05 = *, p<0.01 = **, p<0.001 = ***, p<0.0001 = ****. (A) Left: Schematic of immunization regimen. Mice were injected at week 0 with mosaic-8b RBD-nanoparticles administered as ALD mix3, ALD mix4, SD, or conventionally (bolus injections with adjuvant). At week 4, conventionally immunized mice were given an additional bolus injection of mosaic-8b RBD-nanoparticles plus adjuvant. Right: Colors used to identify immunizations and symbols used to identify matched (filled in square data points; gray shading around name) or mismatched (unfilled square data points; black outline around name) sarbecovirus antigens. Sarbecovirus strain names are colored throughout the figure to indicate clade. (B) ELISA results for serum samples from weeks 12 and 31 after immunization. Geometric means of ED_50_ values for animals in each cohort (symbols with standard deviations indicated by error bars) are connected by colored lines. Mean titers against RBDs from indicated sarbecoviruses were compared pairwise across immunization cohorts by Tukey’s multiple comparison test with the Geisser-Greenhouse correction (as calculated by GraphPad Prism). Titers were significantly higher for ALD mix3 and ALD mix4 compared to conventional mosaic-8b immunization at week 12 (ALD mix3 vs. mosaic-8b p<0.0001, ALD mix4 vs. mosaic-8b p<0.0001) but not at week 31, when the conventional immunization titers were significantly higher (ALD mix3 vs. mosaic-8b p<0.0001, ALD mix4 vs. mosaic-8b p<0.0001). (C) Neutralization potencies for serum samples from weeks 12 and 31 after immunization presented as half-maximal inhibitory dilutions (ID_50_ values) of sera against pseudoviruses from the indicated mismatched coronavirus strains (results for week 16 are shown in Figure S1A). Dashed horizontal lines correspond to the limit of detection. Data for each immunization group are visualized using box and whisker plots, with each data point representing serum from one animal. The boxes display the range between the upper and lower quartiles, with a line denoting the median value. The whiskers extend to minimum and maximum values, excluding any outliers. Significantly higher neutralization titers were found for the following pairwise comparisons: against XBB.1.5 (ALD mix3 vs. mosaic-8b p=0.0014, ALD mix4 vs. mosaic-8b p=0.0032), SARS-1 (ALD mix3 vs. mosaic-8b p=0.0001, ALD mix4 vs. mosaic-8b p<0.0001), BtKy72 (ALD mix4 vs. mosaic-8b p=0.0232), and Khosta-2 (ALD mix4 vs. mosaic-8b p=0.0003) at week 12 and against XBB.1.5 (ALD mix3 vs. SD p=0.0482), SARS-1 (ALD mix3 vs. SD p<0.0001, ALD mix4 vs. SD p<0.0001, SD vs. mosaic-8b p=0.0025), BtKY72 (ALD mix3 vs. SD p=0.0010, ALD mix4 vs. SD p=0.0045, SD vs. mosaic-8b p=0.0004), and Khosta-2 (ALD mix3 vs. SD p=0.0496, ALD mix4 vs. SD p=0.0055) at week 31.

Mean ELISA binding titers (ED_50_ values) for serum Ab binding to sarbecovirus RBDs were significantly higher for ALD mix3 and ALD mix4 when compared to titers for conventional mosaic-8b immunization at week 12 (Figure 2B, left). However, at week 31, titers for both ALD mix3 and ALD mix4 were significantly reduced compared with the conventional mosaic-8b mean ED_50_ values (Figure 2B, right), suggesting lower durability of ALD-coated mosaic-8b nanoparticle immunizations compared with conventional mosaic-8b vaccination.

In vitro neutralization assays were performed for four mismatched pseudovirus strains: SARS-2 XBB.1.5, SARS-CoV (referred to here as SARS-1), BtKY72, and Khosta-2 (Figure 2C). These strains include (i) a hard-to-neutralize SARS-2 variant (XBB.1.5; RBD related to the WA1 RBD by 89% amino acid sequence identity), (ii) a clade 1a sarbecovirus that spilled over into humans (SARS-1; related by 74% sequence identity to the WA1 RBD), and (iii, iv) clade 3 animal sarbecoviruses (BtKY72 and Khosta-2; related by 73% and 68% sequence identities to WA1 RBD, respectively) with human spillover potential whose RBDs are more distantly related to the clade 1a, 1b, and 2 RBDs present on mosaic-8b RBD-nanoparticles (Figure 1C). At week 12, ALD mix3 samples showed significantly higher heterologous neutralization titers when compared to conventional mosaic-8b vaccination (Figure 2C, left panel) against XBB.1.5, SARS-1, and ALD mix4 immunization further showed significantly higher neutralization titers than conventional mosaic-8b vaccination for BtKy72 and Khosta-2 (Figure 2C, left panel). Indeed, the mix3 and mix4 samples exhibited mean neutralization titers between 10^2^ and 10^4^ against all four viruses, whereas conventionally delivered mosaic-8b either showed weak (Khosta-2), undetectable (XBB.1.5), or 5-10–fold reduced mean neutralization titers (SARS-1 and BtKY72) compared with the ALD samples. Although ALD-delivered mosaic-8b continued to elicit significantly higher neutralization titers than conventional mosaic-8b immunization against matched (SARS-2 Beta, WIV1) and mismatched (both ALD mixes for SARS-2 XBB.1.5; ALD mix4 for SARS-2 KP.3 and Khosta-2) viral strains at week 16 (Figure S1A,B), these differences were not significant at week 31 (Figure 2C, right panel), likely due to Ab contraction as observed for the ELISA ED_50_ values (Figure 2B, right). The lower durability of the Ab responses to ALD-delivered mosaic-8b compared with conventional mosaic 8b immunizations could be a result of adjuvant differences (aluminum for ALD; Addavax for conventional). Nevertheless, these results demonstrate that pulsatile release of mosaic-8b RBD-nanoparticles through ALD coating resulted in improved breadth and potency of Ab responses, including the gain of neutralizing responses to heterologous strains, and an enhanced ability to generate broadly neutralizing serum Abs compared with the analogous vaccine delivered conventionally.

Immune imprinting, originally described for influenza virus infections, refers to preferential boosting of Ab responses against epitopes shared with the first strain of a variable immunogen to which an individual was originally exposed.^20,21^ To explore potential effects of imprinting on mosaic-8b RBD-nanoparticle immunizations, we previously compared immune responses to mosaic-8b RBD-nanoparticles in animals with no previous exposure to SARS-CoV-2 antigens (originally naïve animals) to responses in animals that had been pre-vaccinated with COVID-19 vaccines presenting WA1 SARS-CoV-2 spike. This study showed that, in common with the originally naïve animals, pre-vaccinated cohorts also exhibited cross-reactive Ab binding and neutralizing properties.^13^ Here, we addressed whether prior exposure to a COVID-19 vaccine would affect the ability of ALD-coated mosaic-8b to elicit broader Ab responses by comparing ALD mix3 and ALD mix4 to conventional mosaic-8b immunizations in mice pre-vaccinated with two doses of an mRNA-LNP equivalent to Pfizer-BioNTech’s WA1 spike-encoding BNT162b2 vaccine. Twelve weeks after the second mRNA-LNP vaccine dose, pre-vaccinated mice were immunized with either a single dose of ALD mix3, ALD mix4, or SD antigens, or with two doses of mosaic-8b RBD-nanoparticles (Figure 3A). In these pre-vaccinated mice, ALD mix3 and ALD mix4 immunizations elicited significantly higher mean Ab binding titers than either the conventional mosaic-8b or SD immunizations at both weeks 8 and 17 after immunization (23 and 32 weeks, respectively, after the first vaccine dose), with minimal Ab contraction between the two time points (Figure 3B). At week 8 post-immunization, neutralization titers were higher for ALD mix4 when compared to SD against mismatched XBB.1.5 and SARS-1 viruses. At week 12 post-immunization, neutralization titers against SARS-2 D614G (RBD matched to the pre-vaccination strain) and SARS-2 Beta (matched to the SARS-2 Beta RBD on mosaic-8b) were roughly equivalent (mean ID_50_ titers across the four immunization groups of ∼10^4^) (Figure S1C,D). However, significant improvements for ALD-delivered immunizations compared with conventional immunization were found for neutralization of mismatched (XBB.1.5, KP.3, and Khosta-2 strains) and for strains matched to mosaic-8b (WIV1 and SHC014) (Figure S1D). By week 17, ALD mix3 and ALD mix4 showed significantly higher neutralizing titers than mosaic-8b immunization against mismatched strains (both ALD mixes for XBB.1.5 and SARS-1; ALD mix4 for BtKY72, and Khosta-2) (Figure 3C). Thus, pulsatile release of mosaic-8b via ALD coating improved neutralizing Ab responses under conditions of immunological imprinting to the WA1 SARS-2 variant.

**Figure 3.**
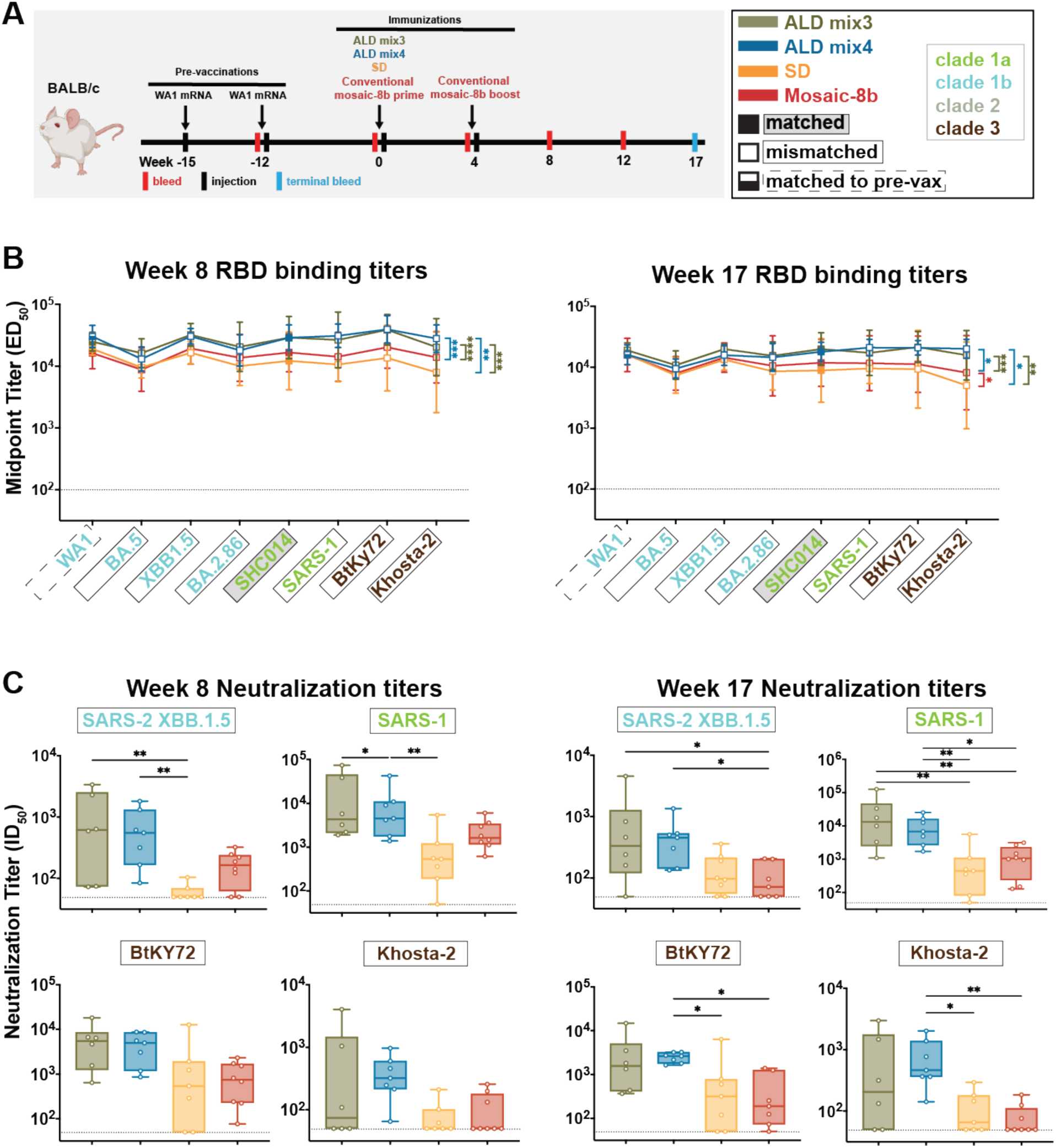
ALD-coated mosaic-8b immunization elicits more broadly neutralizing Ab responses than conventional mosaic-8b immunizations in mice pre-vaccinated with mRNA-LNP encoding WA1 spike. Significant differences between cohorts linked by vertical lines in panels B and C are indicated by asterisks: p<0.05 = *, p<0.01 = **, p<0.001 = ***, p<0.0001 = ****. (A) Left: Schematic of vaccination/immunization regimen. Mice were vaccinated at weeks −15 and −12 (3 weeks apart) with mRNA-LNP vaccines encoding the WA1 spike. At week 0 (15 weeks after the first vaccination), mice were injected with mosaic-8b RBD-nanoparticles administered as ALD mix3, ALD mix4, SD, or conventionally (bolus injection with adjuvant). At week 4, conventionally immunized mice were given an additional bolus injection of mosaic-8b RBD-nanoparticles plus adjuvant. Right: Colors used to identify immunizations and symbols used to identify matched (filled in square data points; gray shading around name), matched to the WA1 pre-vaccination (half-filled in square data points; dashed black outline around name), or mismatched (unfilled square data points; black outline around name) sarbecovirus antigens. Colors used throughout the figure indicate clades of sarbecovirus strains. (B) ELISA results for serum samples from weeks 8 and 17 after immunization. Geometric means of ED_50_ values for animals in each cohort (symbols with standard deviations indicated by error bars) are connected by colored lines. Mean titers against RBDs from indicated sarbecoviruses were compared pairwise across immunization cohorts by Tukey’s multiple comparison test with the Geisser-Greenhouse correction (as calculated by GraphPad Prism). Binding titers for ALD mix3 and ALD mix4 were significantly higher than the conventional mosaic-8b or SD cohorts at weeks 8 (ALD mix3 vs. mosaic-8b p<0.0001, ALD mix4 vs. mosaic-8b p=0.0002) and 17 (ALD mix3 vs. mosaic-8b p<0.0008, ALD mix4 vs. mosaic-8b p=0.0315). (C) Neutralization potencies for serum samples from weeks 8 and 17 after immunization presented as half-maximal inhibitory dilutions (ID_50_ values) of sera against pseudoviruses from the indicated mismatched coronavirus strains (results for week 12 are shown in Figure S1B). Data for each immunization group are visualized using box and whisker plots, with each data point representing serum from one animal. The boxes display the range between the upper and lower quartiles, with a line denoting the median value. The whiskers extend to minimum and maximum values, excluding any outliers. Neutralization titers for ALD mix3 and ALD mix4 were significantly higher than the conventional mosaic-8b or SD cohorts at week 8 against SARS-2 XBB.1.5 (ALD mix3 vs. SD p=0.0042, ALD mix4 vs. SD p=0.0047) and SARS-1 (ALD mix4 vs. SD p=0.0066) and at week 17 against XBB.1.5 (ALD mix3 vs. SD p=0.0433, ALD mix4 vs. SD p=0.0410), SARS-1 (ALD mix3 vs. mosaic-8b p=0.0079, ALD mix4 vs. mosaic-8b p=0.0310, ALD mix3 vs. SD p=0.0010, ALD mix4 vs. SD p=0.0038), BtKY72 (ALD mix4 vs. mosaic-8b p=0.0124, ALD mix4 vs. SD p=0.0317), and Khosta-2 (ALD mix4 vs. mosaic-8b p=0.0029, ALD mix4 vs. SD p=0.0145).

### Epitope mapping using DMS revealed a polyclass Ab signature with ALD-delivered mosaic-8b immunization

We next investigated which RBD epitopes were targeted by Abs elicited by ALD delivery of mosaic-8b RBD-nanoparticles. Ab epitopes on RBDs were previously classified based on structural properties and their degree of conservation across sarbecoviruses, with class 1 and class 2 RBD epitopes exhibiting more sequence variability across sarbecoviruses and SARS-2 variants, and class 4, class 1/4, and portions of class 3 and class 5 epitopes being more conserved^22–25^ (Figure 1E; Figure 4A). In control experiments in which we evaluated equimolar mixtures of anti-RBD monoclonal Abs that targeted multiple epitopes, we observed weak escape profiles with no clear features, which we defined as a “polyclass” response.^13^ Thus, polyclass DMS profiles might be produced in DMS experiments with polyclonal sera containing multiple classes of anti-RBD Abs.

**Figure 4.**
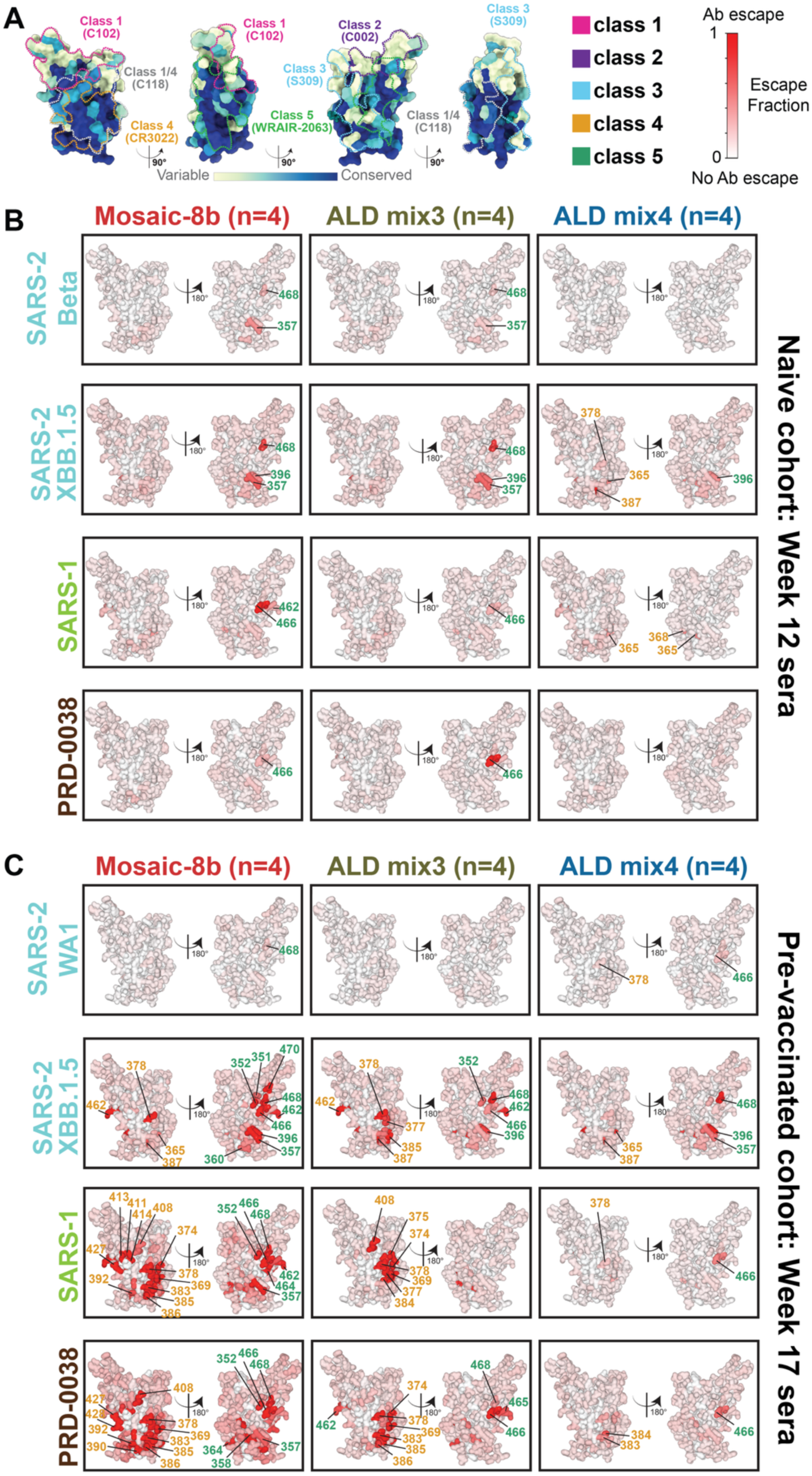
ALD coated mosaic-8b immunization elicits a more polyclass Ab response than conventional mosaic-8b immunization in both naïve and pre-vaccinated mice. (A) ConSurf^44^ calculation of sequence conservation of 16 sarbecovirus RBDs plotted on a surface representation of SARS-2 RBD (PDB 7BZ5). Anti-RBD class 1, 2, 3, 4, 1/4, and 5 Ab epitopes^22–25^ are outlined in dots in different colors using representative structures of Abs bound to SARS-2 spike or RBD (C102: PDB 7K8M; C002: PDB 7K8T, S309: PDB 7JX3; CR3022: PDB 7LOP; C118: PDB 7RKV; WRAIR-2063: PDB 8EOO). (B,C) The average site-total Ab escape calculated for results from n = 4 samples for the RBD libraries indicated on the left. Mice were immunized with the indicated immunogens, and results were mapped to the surface of the WA1 RBD (PDB 6M0J). Gray indicates no escape, and a gradient of red represents the indicated degree of escape. Residue numbers show sites with the most escape with the font colors indicating different RBD epitopes (defined in panel A). The same data are shown in Figure S2 (line plots) and Figures S3, S4 (logo plots) and are summarized in Table 1.

**Table 1.**
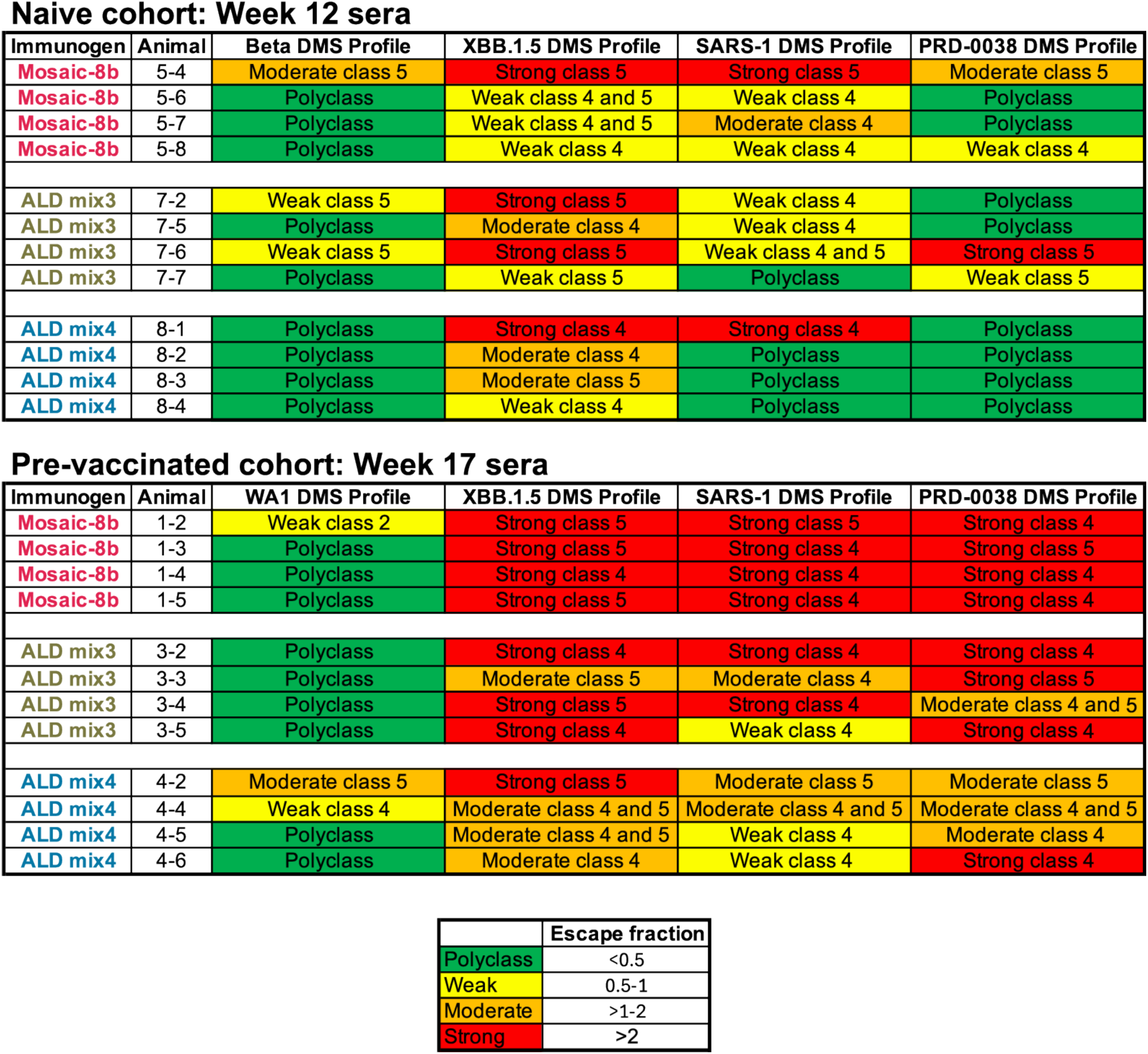
Summary of DMS data (Figures 4, S2, S3, S4) showing that ALD-coated mosaic-8b immunization elicits a more polyclass Ab response than conventional mosaic-8b immunization in both naïve and pre-vaccinated mice. DMS profiles were classified as polyclass, weak, moderate, or strong as indicated by the maximum escape fraction for any RBD position, as indicated in the legend at the bottom.

To characterize epitopes recognized by polyclonal Abs raised after immunization with either ALD and non-ALD antigens, we conducted DMS^16^ using yeast RBD display libraries derived from Beta, WA1, XBB.1.5, SARS-1, and PRD-0038 spikes, which allowed epitope identification by finding residues where substitutions affected binding of polyclonal Abs. We used a SARS-2 Beta library for serum from the naïve and then mosaic-8b immunized animals, and a SARS-2 WA1 library for serum from animals vaccinated with an mRNA-based COVID-19 vaccine and then immunized with mosaic-8b to enable mapping of Ab responses against an RBD matched to the first exposure (SARS-2 Beta is a matched RBD for the originally naïve mice immunized with mosaic-8b; WA1 is a matched RBD for mice pre-vaccinated with WA1 spike-encoding mRNA-LNP). We also used SARS-2 XBB.1.5, SARS-1, and PRD-0038 RBD libraries to map Ab responses to mismatched strains, thereby accounting for DMS results potentially depending on which RBD library was used (e.g., any class 1 and class 2 anti-RBD Abs elicited by mosaic-8b would be unlikely to bind to XBB.1.5, SARS-1, or PRD00-38 RBDs and would therefore not be detected).

For immunizations of the originally naïve cohort, we found that conventional mosaic-8b elicited Abs that recognized class 4 and class 5 RBD epitopes (Figure 4A), as indicated by escape profiles at residues within these epitopes (Table 1, Figure 4B, Figure S2A, S3). Similar results were previously found using a Beta RBD library^14^ and observed here for the XBB.1.5, SARS-1, and PRD-0038 RBD libraries. In contrast, ALD mix4, and to a lesser extent ALD mix3, showed a more polyclass response or a shifted DMS escape profile at residues within class 4 and class 5 RBD epitopes (Table 1, Figure 4B, Figure S2A, S3). Specifically, ALD mix4 showed lower escape fractions than the other two groups against the Beta and PRD-0038 libraries, while escape was shifted to a more class 4 response against XBB.1.5 and SARS-1 libraries as compared to the conventional mosaic-8.

For immunizations of the pre-vaccinated cohort, we found that Ab escape profiles for the SARS-2 WA1 RBD library were predominantly polyclass after mosaic-8b, ALD-mix3, and ALD-mix4 immunizations (Table 1, Figure 4C, Figure S2B, S4), similar to results observed previously for mosaic-8b immunizations in mice that were pre-vaccinated with various COVID-19 vaccines.^13^ In contrast, for the XBB.1.5, SARS-1, and PRD-0038 libraries, conventional mosaic-8b antisera showed strong escape mapping to class 4 and class 5 residues, with ALD mix4 sera showing weak to moderate escape profiles (polyclass with some class 4 and class 5 escape), and ALD mix3 showing an escape profile with characteristics falling between conventional mosaic-8b and ALD mix4 (Table 1, Figure 4C, Figure S2B, Figure S4).

### ALD-delivered mosaic-8b RBD-nanoparticles elicited IgG1-dominated Ab responses

IgG subclasses can elicit distinct Fc effector functions including opsonization, phagocytosis, and Ab-dependent cell-mediated cytotoxicity, which are mediated by differential binding of their Fc domains to Fc gamma receptors (FcγRs).^26^ For example, mouse IgG1, IgG2a, and IgG2b bind with µM affinities to FcγR2b (an inhibitory receptor) and to FcγR3 (an activating receptor), whereas only IgG2a and IgG2b bind to FcγR4 (an activating receptor), each with 200-300 nM affinities, and mouse IgG3 does not bind detectably to any of these FcγRs.^27^ Of the two mouse activating receptors, FcγR3 is more widely expressed than FcγR4, with FcγR4 being mainly found only on monocytes/macrophages and neutrophils and FcγR3 found on those cells plus on dendritic cells, natural killer cells, basophils, mast cells, and eosinophils.^27^

We used systems serology^28^ to evaluate the distributions of IgG subclasses and FcγR-binding IgGs elicited by mosaic-8b RBD-nanoparticles delivered either conventionally or by ALD. In these experiments, we assessed binding of total IgG versus IgG1, IgG2a, IgG2b, IgG3, and FcγR2b-, FcγR3-, or FcγR4-binding IgGs to a panel of spike trimers and RBDs derived from SARS-2 variants and other sarbecoviruses (Figure 5).

**Figure 5.**
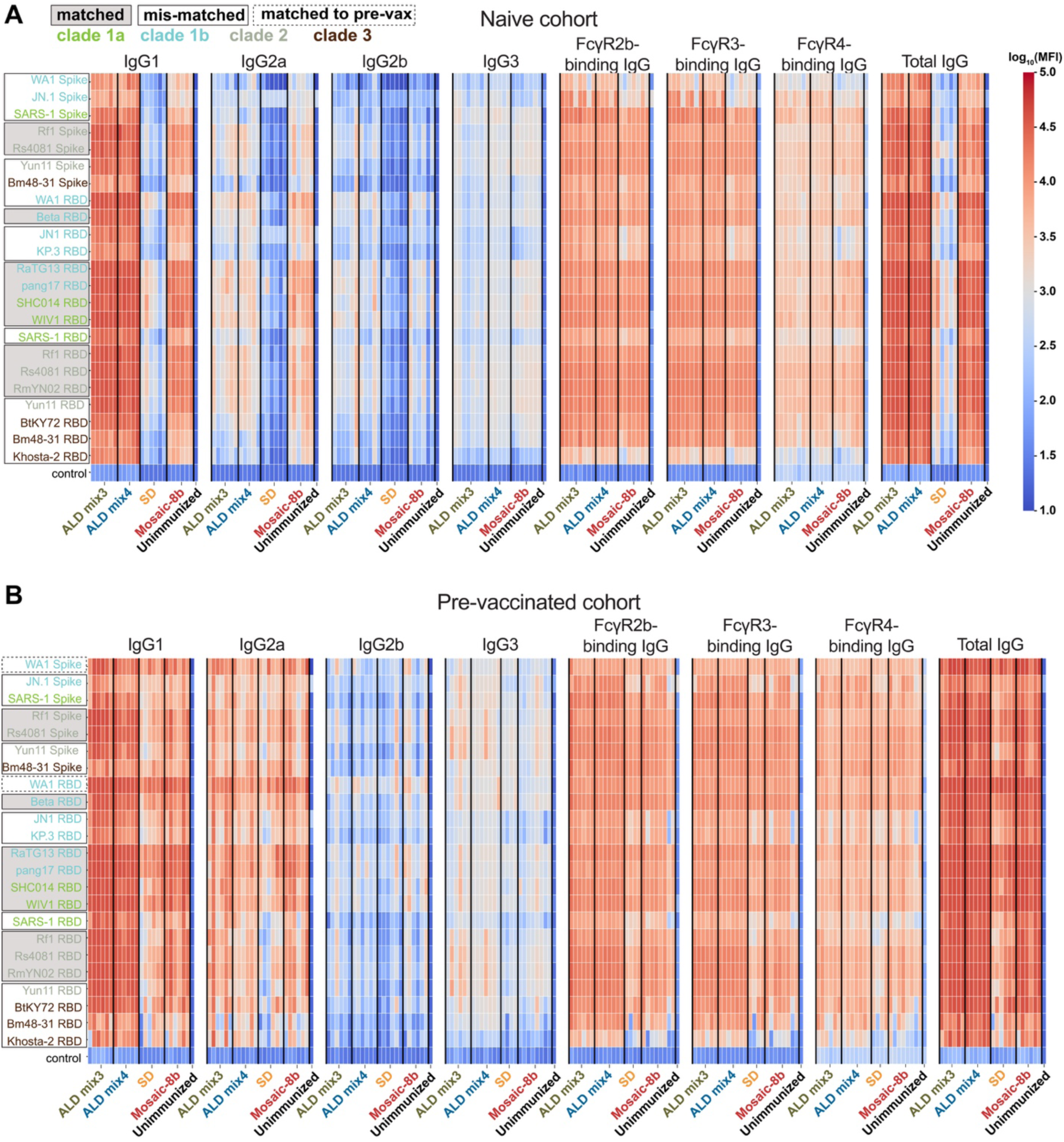
ALD-coated mosaic-8b immunization elicits an IgG1-dominated response. Analysis of binding interactions of IgG1, IgG2a, IgG2b, IgG3, FcγR2b-binding IgGs, FcγR3-binding IgGs, FcγR4-binding IgGs, and total IgG with the indicated spikes, RBDs, or a non-sarbecovirus control protein. Antigen names (y-axes) are colored to identify clades from which spikes or RBDs were derived. Matched antigens are indicated with gray shading around the name, mismatched antigens are indicated with a black outline around the name, and WA1 antigens that are matched to the WA1 pre-vaccination are indicated with a dashed black outline around the name. Serum from mice immunized with ALD mix3, ALD mix4, SD, and conventional mosaic-8b (x-axes) in originally naïve (A) or pre-vaccinated (B) cohorts was tested for binding against different sarbecovirus spikes and RBDs (y-axes). Each vertical line of binding data represents an individual mouse and immunization groups are separated by a vertical black line. Unimmunized = Day 0 serum from a naïve mouse prior to mosaic-8b immunization. MFI = median fluorescent intensity.

In originally naïve animals that were then immunized with the differently-delivered mosaic-8b RBD-nanoparticles (Figure 2A), IgG1 titers for binding to a panel of sarbecovirus RBDs were significantly higher for the ALD mix3 and mix4 groups compared to the other groups, and all groups showed lower titers for IgG2a, IgG2b, and IgG3 than for IgG1 (Figure 5A, Figure 6A).

**Figure 6.**
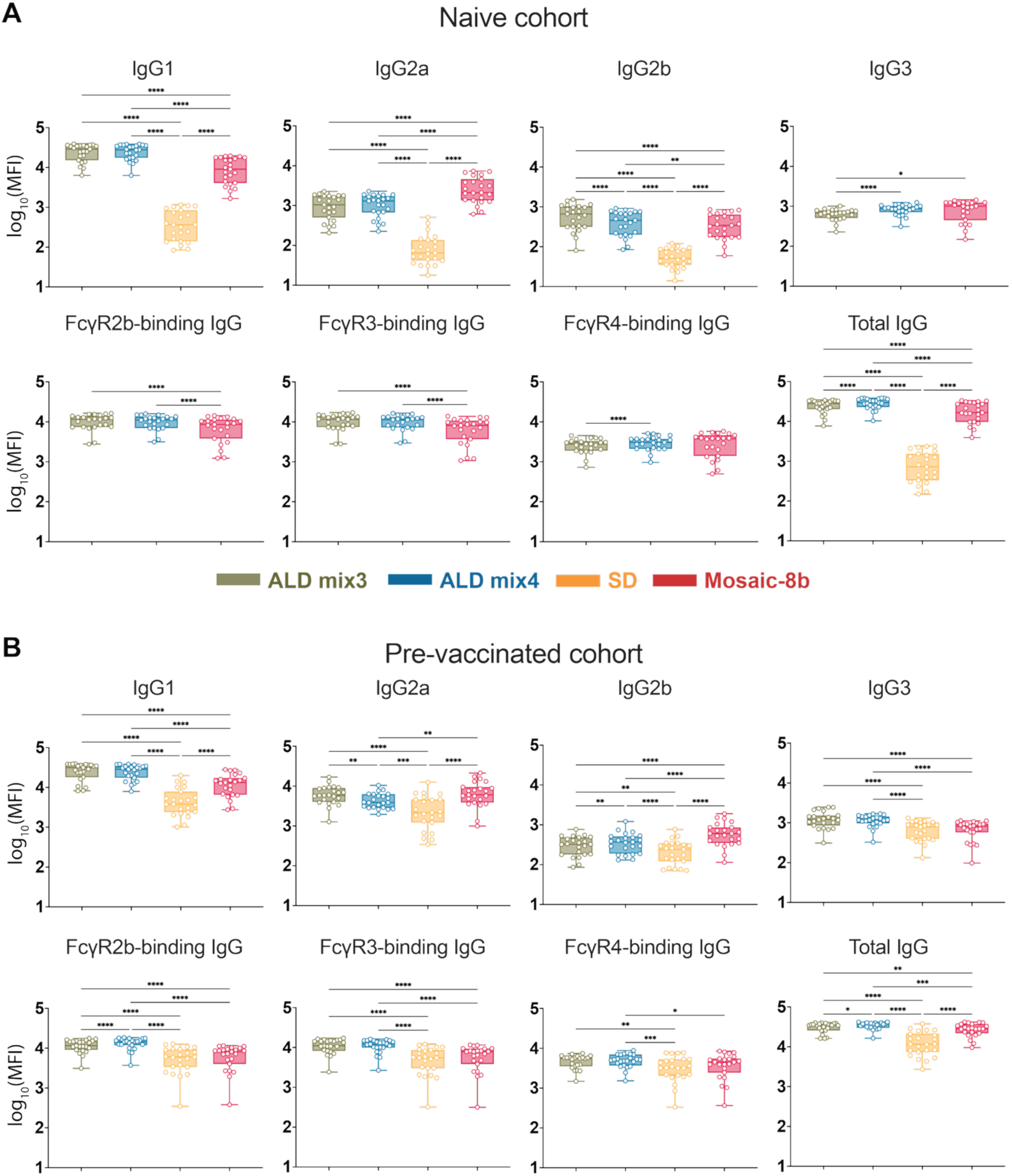
ALD-coated mosaic-8b immunization elicits different IgG subclass and FcγR responses than conventional mosaic-8b immunizations. For IgG1, IgG2a, IgG2b, IgG3, FcγR2b-binding IgGs, FcγR3-binding IgGs, FcγR4-binding IgGs, and total IgG, geomean MFI (geometric median fluorescent intensity) values from the naïve cohort (A) or the pre-vaccinated cohort (B) are represented as points corresponding to different spikes or RBDs (individual responses shown in Figure 5) and compared pairwise across immunization cohorts by Tukey’s multiple comparison test calculated by GraphPad Prism. Significant differences between cohorts linked by vertical lines in panels B and C are indicated by asterisks: p<0.05 = *, p<0.01 = **, p<0.001 = ***, p<0.0001 = ****.

This result might be related to the microparticulate nature of the alumina-coated vaccines, which could function to stimulate cellular uptake and act as an alum-like adjuvant to stimulate IgG1-dominated TH2 responses in mice.^29,30^ In contrast, IgG2a responses were significantly higher for conventional mosaic-8b, whereas ALD mix3 and ALD mix 4 elicited significantly higher IgG2b titers than conventionally delivered mosaic-8b (Figure 5A, Figure 6A). Consistent with IgG1 titer levels, ALD mix3 and ALD mix4 elicited significantly higher FcγR2b-, and FcγR3-binding IgGs (Figure 5A, Figure 6A), but lower levels of antigen-specific FcγR4-binding IgGs (Figure 5A), consistent with mouse FcγR4 specificity for IgG2a and IgG2b but not IgG1.^27^ There was no difference in overall FcγR4-binding IgGs when comparing ALD mix3 and ALD mix4 to conventionally delivered mosaic-8b, although the FcγR4-binding IgG responses were significantly higher for ALD mix4 when compared to ALD mix3. Finally, as expected, binding responses of total IgG (Figure 5A) were consistent with ELISA binding titers (Figure 2B) showing broad recognition across sarbecovirus RBDs, with significantly higher titers elicited by ALD mix3 and ALD mix4 compared with the other groups. Together, this suggests that ALD mix3 and ALD mix4 elicited equivalent or stronger Ab responses capable of eliciting Fc effector functions compared with conventional mosaic-8b delivery.

In the pre-vaccinated animals, IgG1 dominated the ALD mix3 and ALD mix4 responses, although IgG2a responses in all groups were higher than for the naïve cohort (Figure 5B, Figure 6B), likely due to recall of mRNA LNP-elicited IgG2a class-switched B cells.^31^ IgG2a and IgG2b responses were significantly higher for conventionally delivered mosaic-8b than for ALD mix3 and ALD mix4. Similar to the naïve cohort, FcγR2b- and FcγR3-binding IgGs were significantly higher for ALD mix3 and ALD mix4 than for conventionally delivered mosaic-8b (Figure 5B, Figure 6B). ALD mix4 elicited FcγR4-binding IgGs were also significantly higher than for conventional mosaic-8b (Figure 5B, Figure 6B). Again, total IgG binding responses were also broad across RBDs for all four cohorts (Figure 5B), consistent with ELISAs (Figure 3B). Together, these results show that ALD-delivered mosaic-8b maintained a stronger Ab response capable of eliciting broad Fc effector functions as compared with conventionally delivered mosaic-8b in both previously vaccinated and originally naïve animals.

## Discussion

Vaccines capable of eliciting broadly cross-reactive Abs that recognize conserved motifs expressed across variants of a microbe are desirable for a broad range of infectious diseases.^32^ Here, we show that a single injection of a candidate pan-sarbecovirus vaccine delivered within microparticles coated with alumina by ALD increases the breadth of elicited Abs and changes the distribution of induced IgG subclasses and FcγR binding properties compared with typical prime-boost bolus injections of the same immunogen. Furthermore, Ab binding titers generated by mosaic-8b RBD-nanoparticles embedded within ALD-coated microparticles were higher than those generated by conventional formulations, consistent with reports for vaccines using other antigens.^4,6,30^ Technologies such as ALD that enable exposure of related antigens over time^4–6^ to guide elicitation of broadly neutralizing Abs will be critical for the practical success of a vaccine approach that requires a sequential series of immunogens (e.g., as is likely required for a successful HIV-1 vaccine^33^).

Here, we investigated whether immune responses elicited by pulsatile delivery of a single immunogen, mosaic-8b RBD-nanoparticles, would be distinct from responses elicited by conventional prime-boost bolus immunizations of the same immunogen. Mosaic-8b presents eight different SpyTagged sarbecovirus RBDs coupled randomly to the 60 attachment sites on a SpyCatcher-mi3 nanoparticle.^19^ Conventional immunizations of mosaic-8b RBD-nanoparticles induce broad Ab responses against zoonotic sarbecoviruses and SARS-CoV-2 variants of concern in mice,^13–15,34^ non-human primates,^14^ and rabbits,^35^ and increased protection against a mismatched challenge (i.e., a virus not represented by an RBD on mosaic-8b) compared with homotypic (only SARS-CoV-2 RBDs) RBD-nanoparticles.^14^ The question we address here is whether Ab responses induced by ALD delivery of mosaic-8b show improvements and/or differences compared with conventional mosaic-8b immunizations, which is of relevance to vaccine protocols in general and to this vaccine candidate in particular. To mimic potential use of mosaic-8b RBD-nanoparticles in humans, most of whom have been exposed to SARS-CoV-2 through vaccination, infection, or both, we compared results from mice that were originally naïve to SARS-2 antigens prior to mosaic-8b immunization and from mice that had been pre-vaccinated with a prime-boost mRNA-based vaccine series prior to mosaic-8b immunization.

We found that two versions of ALD-formulated mosaic-8b (ALD mix3 and ALD mix4) both induced the broad Ab binding and neutralization properties observed for conventional mosaic-8b immunizations in mice and other animal models.^14,35^ We found significantly higher mean binding titers from the ALD cohorts across a panel of eight sarbecovirus RBDs (seven of which were mismatched with respect to mosaic-8b) at week 12 after initial mosaic-8b immunizations compared with conventional mosaic-8b titers in mice that were originally naïve to sarbecovirus antigens. By week 31, however, mean binding titers for conventionally delivered mosaic-8b were significantly higher than titers induced by ALD delivery. The apparently decreased durability of the ALD-induced versus conventional delivery-induced responses to mosaic-8b might be improved by including an adjuvant. In fact, earlier studies of vaccines using human papilloma virus capsomere antigens formulated together with the aluminum hydroxide adjuvant Alhydrogel^®^ inside ALD-coated microparticles showed no decreases in binding titers at the end of an 11-week study.^6^ In addition, in contrast to the current study, IgG1 titers for an ALD-coated ovalbumin immunization and for a conventional immunization adjuvanted with the alumina-based Alhydrogel^®^ showed similar rates of loss over time.^30^ In our studies, we note that mice that had been pre-vaccinated with a prime-boost series of an mRNA-based COVID-19 vaccine, ALD mix3- and ALD mix4-induced binding responses remained higher than responses induced by conventional mosaic-8b delivery at both weeks 8 and 17 after initial mosaic-8b immunizations.

Demonstrating promise for using ALD to deliver mosaic-8b and potentially also vaccines against mutating pathogens in general, we found that ALD mix3 and ALD mix4 induced Abs that showed increased abilities to neutralize mismatched viruses. For example, week 12 serum from the originally naïve ALD cohorts neutralized the SARS-2 variant XBB.1.5^36^ (15 of 16 mice in ALD mix3 and ALD mix4 cohorts with neutralization titers well above background), whereas week 12 serum from all mice in the conventionally immunized cohort (a time point after both prime and boost bolus immunizations) showed no detectable neutralization of XBB.1.5, whose RBD is related by 90% to the SARS-2 Beta RBD on mosaic-8b). ALD mix4 also showed improved neutralization of SARS-2 KP.3 at week 16 (the KP.3 RBD is related by 86% sequence identity to the SARS-2 Beta RBD on mosaic-8b). By week 31, neutralization titers from the ALD cohort sera had contracted such that only four of eight mice in the ALD mix3 cohort showed ID_50_ values above 10^2^ against XBB.1.5. At this time point, however, only one serum sample from the conventionally immunized cohort had an XBB.1.5 titer above 10^2^. Thus, ALD delivery of mosaic-8b improved elicited Ab neutralization of representative SARS-2 variants of concern. Neutralization titers against other mismatched sarbecoviruses (SARS-1, BtKY72, Khosta-2) also showed improvements for serum samples from the ALD cohorts. In these experiments, SARS-1 (a clade 1a sarbecovirus) represents an as-yet-unidentified virus that could spill over into humans, and BtKY72 and Khosta-2 are distantly-related clade 3 sarbecoviruses with RBDs sharing 62-75% (BtKY72) and 58-72% (Khosta-2) sequence identities with the clade 1a, clade 1b, and clade 2 RBDs present on mosaic-8b, again supporting the use of ALD delivery for mosaic-8b immunizations to elicit more broadly neutralizing Abs. Of relevance to humans with pre-existing immunity to SARS-2 antigens, conventionally delivered mosaic-8b in pre-vaccinated mice elicited higher neutralization titers than the counterpart immunizations in the originally naïve mice against the four mismatched sarbecoviruses, whereas ALD-delivered mosaic-8b elicited superior neutralization titers in nearly all mice. Taken together, the serum Ab binding and neutralization results suggest that ALD delivery of mosaic-8b RBD-nanoparticles would increase its already broadly cross-reactive Ab-eliciting properties, and as such, ALD technology could increase cross-reactive responses elicited by vaccines against other mutating viruses, e.g., influenza and HIV-1.

Using DMS, we mapped the Ab epitopes targeted on four different RBDs in samples from the naïve cohort, one matched (SARS-2 Beta) and three mismatched (XBB.1.5, SARS-1, and PRD-0038), allowing identification of which epitope(s) are associated with better neutralization. ALD-delivered mosaic-8b, particularly ALD mix4, elicited a more polyclass profile (i.e., no clear features in DMS the DMS escape profile^13^) against three of the four strains (all except XBB.1.5) compared to conventionally delivered mosaic-8b. When escape was seen for ALD mix4 antisera, it was mostly represented by class 4 RBD responses (three of four mice for XBB.1.5 and one of four for SARS-1) with less class 5 (one of four for XBB.1.5), whereas conventionally delivered mosaic-8b elicited a mix of class 4 (three of four for XBB.1.5, three of four for SARS-1, one of four for PRD-0038) and class 5 (one of four for Beta, three of four for XBB.1.1.5, one of four for SARS-1, and one of four for PRD-0038). Altogether, the diminished escape elicited by ALD formulations of mosaic-8b suggested that they induced a more evenly distributed and/or less escapable anti-RBD response than conventionally delivered bolus injections of mosaic-8b. This more evenly distributed, or polyclass DMS response, could result from elicitation of Abs that recognize different RBD epitopes,^13^ i.e., in this case, an equal mix of class 4 and class 5 anti-RBD Abs. Alternatively, or in combination with inducing polyclass Abs, ALD-coated mosaic-8b immunizations could induce higher affinity Abs that are more resistant to single RBD substitutions represented in the DMS libraries. The differences in neutralization potency elicited by ALD mix3 and ALD mix4 versus conventionally delivered mosaic-8b does not appear to be because of differential epitope targeting (all groups targeted polyclass or a mix of conserved class 4 and class 5 epitopes) but could rather depend on induction of either higher overall titers of Abs or higher affinity Abs targeting the same epitopes.

In the pre-vaccinated cohort, ALD-delivered mosaic-8b also elicited weaker escape or a more polyclass response compared to conventionally delivered mosaic-8b, again suggesting that the characteristics of the Abs are different between the groups. For conventionally delivered mosaic-8b the escape profile against XBB.1.5, SARS-1, and PRD-0038 showed stronger class 4 and class 5 escape than for mosaic-8b in the naïve cohort, likely due to the recall of mRNA LNP-elicited Abs in mosaic-8b boosted animals.^13^

We also found intriguing differences in subclass and FcγR-binding properties for IgGs elicited by mosaic-8b RBD-nanoparticles delivered as conventional prime-boost bolus injections versus as pulsatile release from a single ALD immunization. Using systems serology,^28^ we showed for the originally naïve cohort that ALD mix3 and ALD mix4 elicited an IgG1-skewed Ab response that exhibited significantly higher FcγR2b- and FcγR3-binding IgGs than the conventional prime-boost bolus immunizations, whereas conventional mosaic-8b immunizations elicited significantly higher IgG2a binding responses. However, ALD mix3, ALD mix4, and conventional mosaic-8b immunizations exhibited similar levels of IgG2b, IgG3, and FcγR4-binding IgGs. This suggests that ALD-delivered mosaic-8b elicits Abs that can mediate stronger Fc effector functions (a correlate of protection against SARS-2^37–39^) against a wide panel of sarbecovirus strains. Similar to the naïve cohort results, ALD mix3 and ALD mix4 in pre-vaccinated animals showed significantly higher antigen-specific IgG1, and FcγR2b- and FcγR3-binding IgGs. By contrast, in the pre-vaccinated mice, the ALD mixes showed equivalent or sometimes lower IgG2a and IgG2b titers than conventional mosaic-8b, with ALD-mix4 exhibiting significantly higher FcγR4-binding IgG titers. Interestingly, ALD mix3 and ALD mix4 immunization in pre-vaccinated mice induced significantly higher IgG3 levels. Mouse IgG3, in common with human IgG4 that increases in humans who received multiple COVID-19 vaccinations,^40^ is a non-inflammatory IgG that does not bind detectably to FcγRs.^41^. This could be related to the multiple pulses of antigen release for the ALD mix3 and ALD mix4 groups. Altogether, these results show that ALD mix3 and ALD mix4 exhibited high FcγR-binding IgG responses, especially in the pre-vaccinated cohort. These responses could likely be further improved by additional use of a Th1-inducing adjuvant that would boost IgG2a responses (e.g., adding a TLR4 agonist such as MPLA or a TLR7/8 agonist such as imiquimod^42^), and therefore higher FcγR4-binding responses, and perhaps improve durability of ALD-formulated vaccine responses.

In summary, ALD delivery of a vaccine candidate, mosaic-8b RBD-nanoparticles, improved elicitation of the broadly cross-reactive responses observed for this immunogen from conventional prime-boost bolus immunizations. These results support evaluation of ALD technology to deliver vaccines against other rapidly mutating pathogens when broadly cross-reactive Abs are essential for efficacy.

### Limitations of the study

We could not include more recent SARS-2 variants of concern in assays due to limited quantities of mouse serum after initial experiments were performed. In addition, our assays to evaluate immune responses included serum Ab binding and neutralization but not a challenge experiment to assess protection. However, we previously demonstrated that conventionally delivered mosaic-8b RBD-nanoparticles protected from matched and mismatched sarbecovirus challenges in K18-hACE2 transgenic mice,^14^ thus, the superior elicited Ab responses by ALD-delivered mosaic-8b suggest protection would also be observed for the ALD mode of immunogen delivery. For studies in pre-vaccinated mice, we used an mRNA-LNP formulation that may not perform identically to clinically available vaccines because we were unable to obtain licensed Pfizer-BioNTech or Moderna mRNA-LNP vaccines from the respective companies for research purposes. Finally, since IgG subclasses and FcγRs show differences between humans and mice,^27^ conclusions from our systems serology studies may not directly translate to humans.

**Figure S1.**
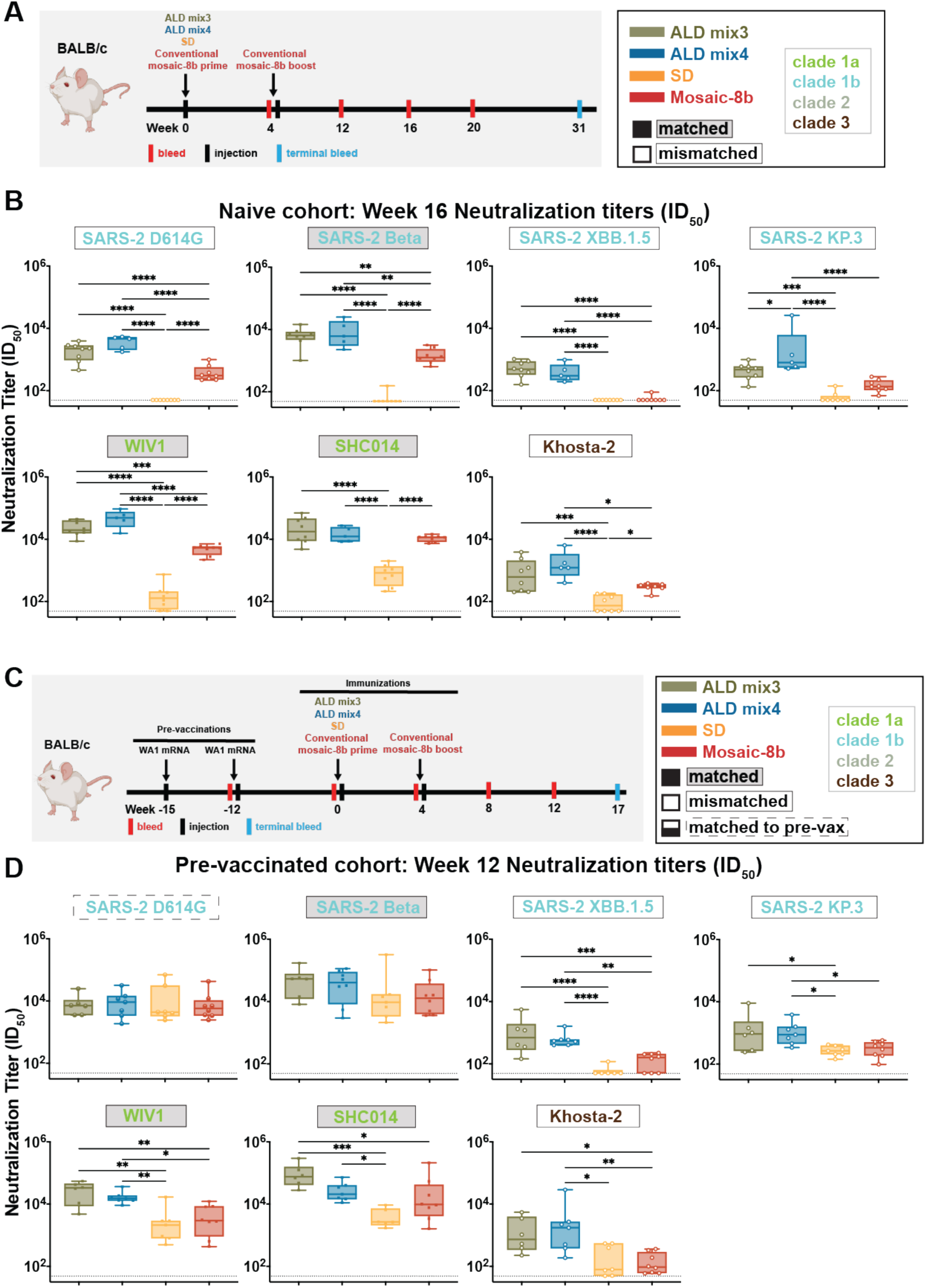
ALD-coated mosaic-8b immunization elicits more broadly matched and mismatched neutralizing Ab responses than conventional mosaic-8b immunizations in naïve and pre-vaccinated mice. Significant differences between cohorts linked by vertical lines in panels B and C are indicated by asterisks: p<0.05 = *, p<0.01 = **, p<0.001 = ***, p<0.0001 = ****. (A) Left: Schematic of immunization regimen for originally naïve mice. Mice were injected at week 0 with mosaic-8b RBD-nanoparticles administered as ALD mix3, ALD mix4, SD, or conventionally (bolus injection with adjuvant). At week 4, conventionally immunized mice were given an additional bolus injection of mosaic-8b RBD-nanoparticles plus adjuvant. Right: Colors used to identify immunizations and symbols used to identify matched (filled in square data points; gray shading around name) or mismatched (unfilled square data points; black outline around name) sarbecovirus antigens. Colors used throughout the figure indicate clades of sarbecovirus strains. (B) Neutralization potencies for serum samples from week 16 after immunization presented as half-maximal inhibitory dilutions (ID_50_ values) of sera against pseudoviruses from the indicated coronavirus strains (results for weeks 12 and 31 are shown in Figure 2). Dashed horizontal lines correspond to the limit of detection. Data for each immunization group are visualized using box and whisker plots, with each data point representing serum from one animal. The boxes display the range between the upper and lower quartiles, with a line denoting the median value. The whiskers extend to minimum and maximum values, excluding any outliers. Significantly higher neutralization titers were found for the following pairwise comparisons: against SARS-2 D614G (ALD mix3 vs. Mosaic-8b p<0.0001, ALD mix3 vs. SD p<0.0001, ALD mix4 vs. Mosaic-8b p<0.0001, ALD mix4 vs. SD p<0.0001), SARS-2 Beta (ALD mix3 vs. Mosaic-8b p=0.0014, ALD mix3 vs. SD p<0.0001, ALD mix4 vs. Mosaic-8b 8b p=0.0011, ALD mix4 vs. SD p<0.0001), SARS-2 XBB.1.5 (ALD mix3 vs. Mosaic-8b p<0.0001, ALD mix3 vs. SD p<0.0001, ALD mix4 vs. Mosaic-8b p<0.0001, ALD mix4 vs. SD p<0.0001), SARS-2 KP.3 (ALD mix3 vs. SD p=0.0003. ALD mix3 vs. ALD mix4 p=0.0326, ALD mix4 vs. Mosaic-8b p<0.0001, ALD mix4 vs. SD p<0.0001), WIV1 (ALD mix3 vs. Mosaic-8b p=0.0004, ALD mix3 vs. SD p<0.0001, ALD mix4 vs. Mosaic-8b p<0.0001, ALD mix4 vs. SD p<0.0001), SHC014 (ALD mix3 vs. SD p<0.0001, ALD mix4 vs. SD p<0.0001), and Khosta-2 (ALD mix3 vs. SD p=0.0003, ALD mix4 vs. Mosaic-8b p=0.0145, ALD mix4 vs. SD p<0.0001) (C) Left: Schematic of vaccination/immunization regimen for pre-vaccinated mice. Mice were vaccinated twice (3 weeks apart) with mRNA-LNP vaccines encoding the WA1 spike. At week 0 (15 weeks after the first vaccination), mice were injected with mosaic-8b RBD-nanoparticles administered as ALD mix3, ALD mix4, SD, or conventionally (bolus injection with adjuvant). At week 4, conventionally immunized mice were given an additional bolus injection of mosaic-8b RBD-nanoparticles plus adjuvant. Right: Right: Colors used to identify immunizations and symbols used to identify matched (filled in square data points; gray shading around name), matched to the WA1 pre-vaccination (half-filled in square data points; dashed black outline around name), or mismatched (unfilled square data points; black outline around name) sarbecovirus antigens. (D) Neutralization potencies for serum samples from week 12 after immunization presented as half-maximal inhibitory dilutions (ID_50_ values) of sera against pseudoviruses from the indicated coronavirus strains (results for weeks 8 and 17 are shown in Figure 3). Data for each immunization group are visualized using box and whisker plots, with each data point representing serum from one animal. The boxes display the range between the upper and lower quartiles, with a line denoting the median value. The whiskers extend to minimum and maximum values, excluding any outliers. Significantly higher neutralization titers were found for the following pairwise combinations: SARS-2 XBB.1.5 (ALD mix3 vs. Mosaic-8b p=0.0008, ALD mix3 vs. SD p<0.0001, ALD mix4 vs. Mosaic-8b p=0.0022, ALD mix4 vs. SD p<0.0001), SARS-2 KP.3 (ALD mix3 vs. SD p=0.0411, ALD mix4 vs. Mosaic-8b p=0.0474, ALD mix4 vs. SD p=0.0314), WIV1 (ALD mix3 vs. Mosaic-8b p=0.0048, ALD mix3 vs. SD p=0.0010, ALD mix4 vs. Mosaic-8b p=0.0159, ALD mix4 vs. SD p=0.0033), SHC014 (ALD mix3 vs. Mosaic-8b p=0.0172, ALD mix3 vs. SD p=0.001, ALD mix4 vs. SD p=0.0123), and Khosta-2 (ALD mix3 vs. Mosaic-8b p=0.0203, ALD mix4 vs. Mosaic-8b p=0.0039, ALD mix4 vs. SD p=0.0115).

**Figure S2.**
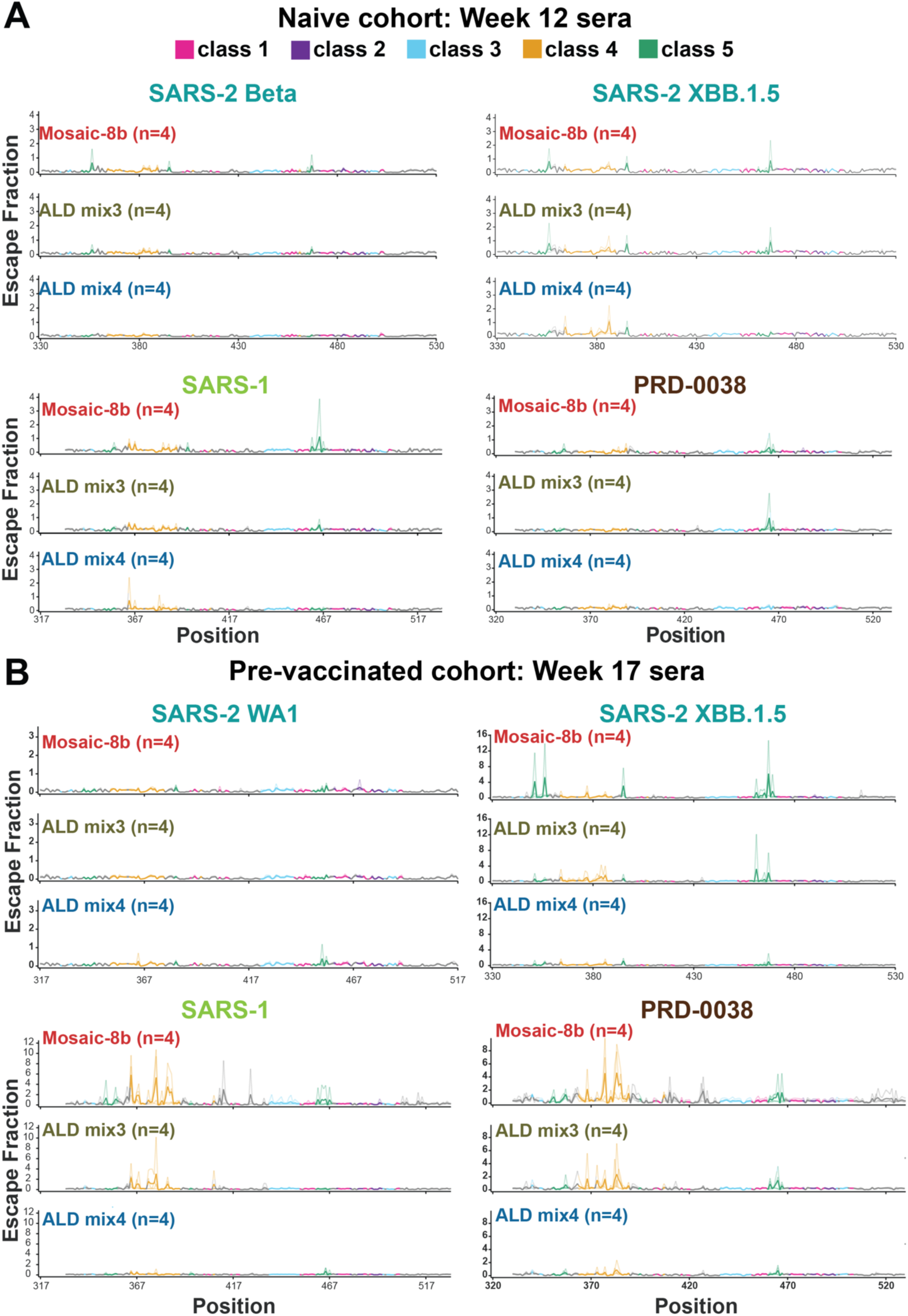
DMS line plots (same data are plotted on RBD structures in Figure 4). (A,B) Line plots for DMS results from the indicated number of samples for the RBD libraries listed at the top of each set of three plots. Mice were immunized with the immunogens indicated above each line plot. X-axis: RBD residue number. Y-axis: sum of the Ab escape of all mutations at a site (larger numbers = more Ab escape). Each line represents one antiserum with heavy lines showing the average across the n=4 sera in each group. Lines are colored according to RBD epitopes in Figure 4A.

**Figure S3.**
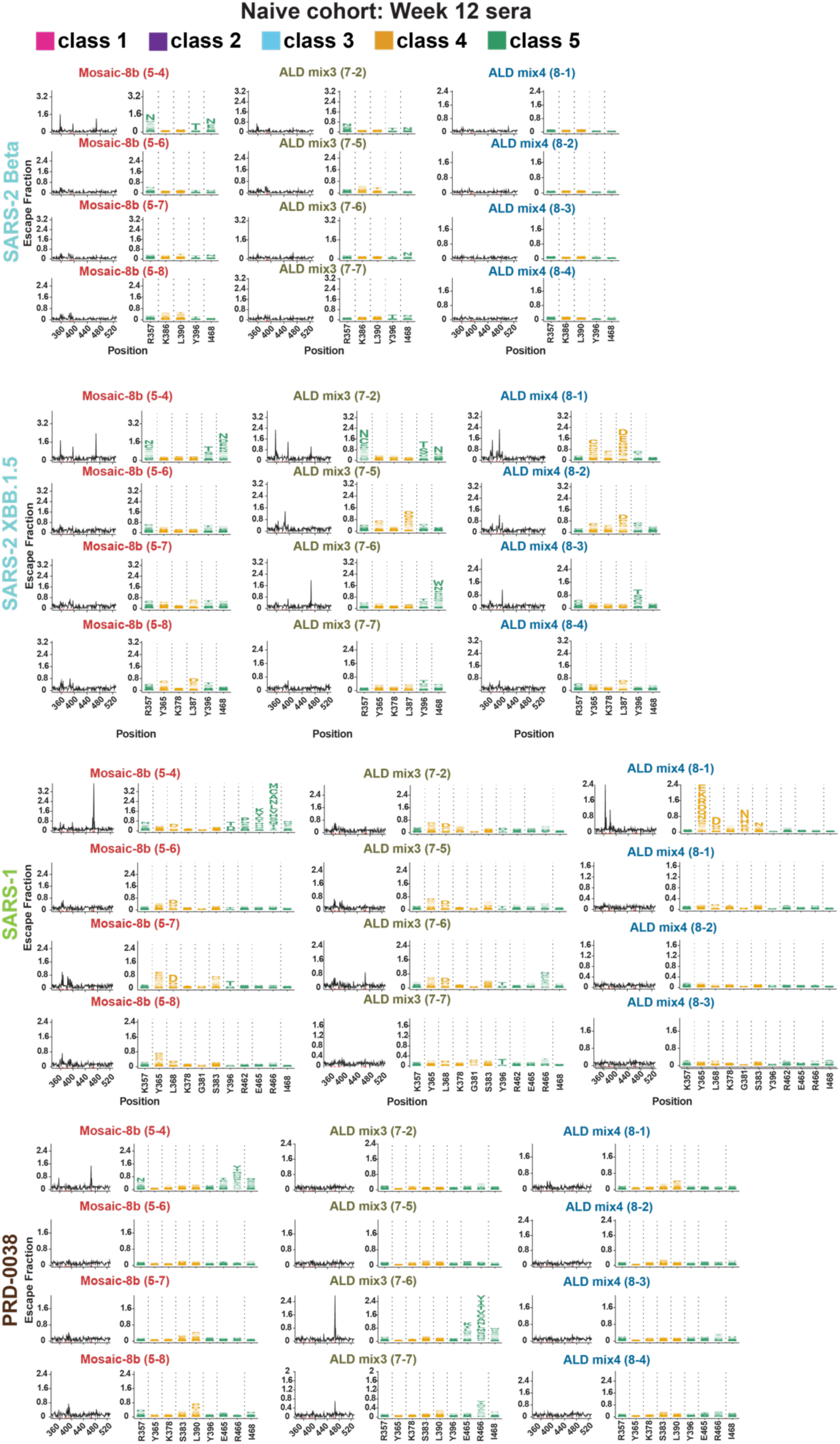
DMS line and logo plots for individual originally naïve mice. Line (left) and logo plots (right) are shown for DMS results for sera from individual mice (IDs in parentheses) collected at 12 weeks after immunization with either mosaic-8b, ALD mix3, or ALD mix4 (immunization schedule in Figure 2A). DMS was performed using the indicated RBD libraries. X-axes of line and logo plots show RBD residue numbers. Y-axes of line plots show the sum of the Ab escape of all mutations at a site (larger numbers indicate increased Ab escape). Sites with the strongest Ab escape are shown in logo plots, with tall letters representing the most frequent mutations at a site. Logo plots are colored for RBD epitopes within different classes^22–25^ (class 1 = pink; class 2 = purple; class 3 = blue; class 4 = yellow; class 5 = green; epitopes are shown in Figures 1E and 4A; gray for residues not assigned to an epitope). Compiled data from all mice in each group are plotted on RBD structures in Figure 4 and as line plots in Figure S2.

**Figure S4.**
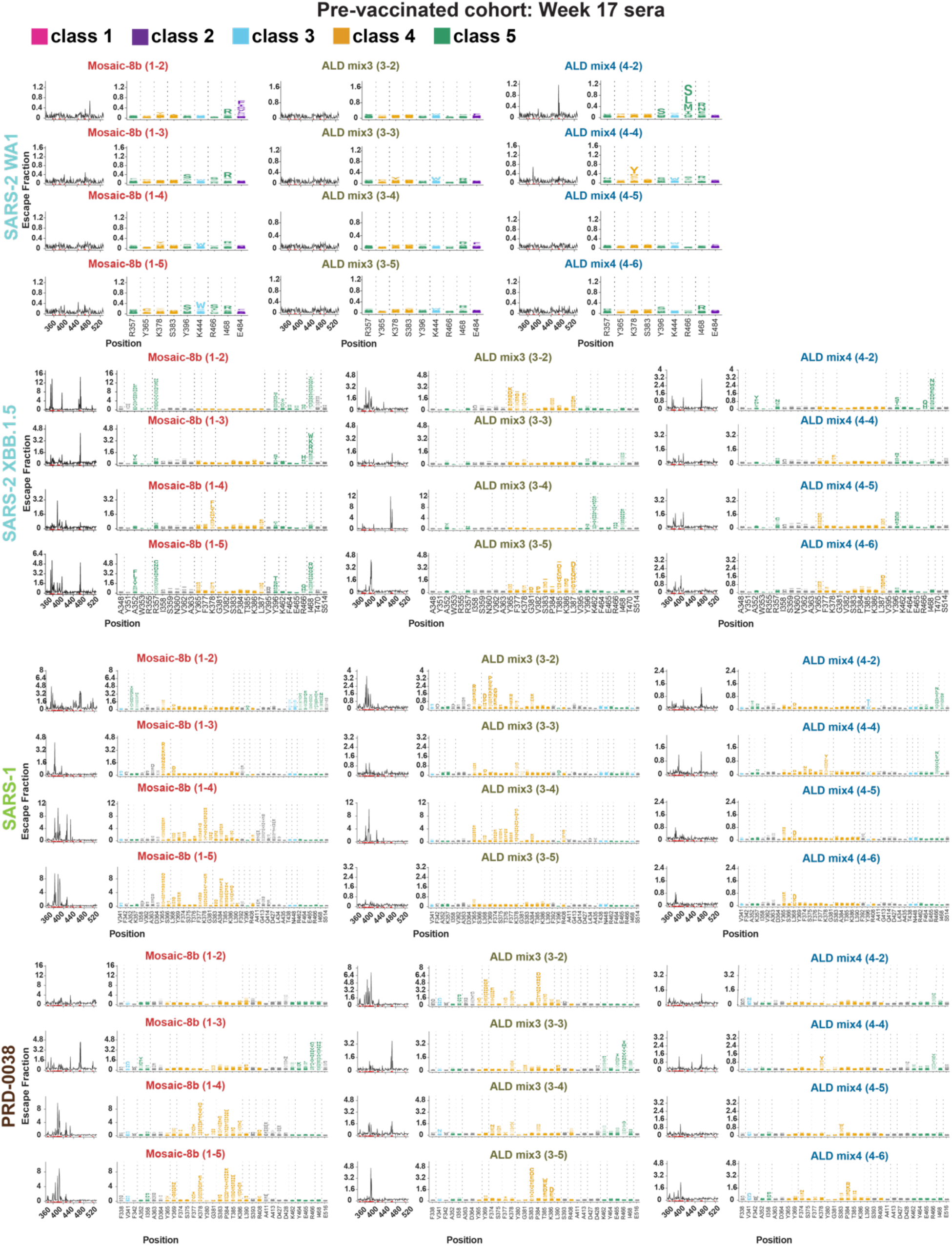
DMS line and logo plots for individual pre-vaccinated mice. Line (left) and logo plots (right) are shown for DMS results at week 17 for sera from individual mice (IDs in parentheses) that were pre-vaccinated with two doses of WA1 mRNA-LNP and then immunized with either mosaic-8b, ALD mix3, or ALD mix4 (vaccination/immunization schedule in Figure 3A). DMS was performed using the indicated RBD libraries. X-axes of line and logo plots show RBD residue numbers. Y-axes of line plots show the sum of the Ab escape of all mutations at a site (larger numbers indicate increased Ab escape). Sites with the strongest Ab escape are shown in logo plots, with tall letters representing the most frequent mutations at a site. Logo plots are colored differently for RBD epitopes within different classes^22–25^ (class 1 = pink; class 2 = purple; class 3 = blue; class 4 = yellow; class 5 = green; epitopes are shown in Figures 1E and 4A; gray for residues not assigned to an epitope). Compiled data from all mice in each group are plotted on RBD structures in Figure 4 and as line plots in Figure S2.

## METHODS

### Preparation of spike and RBD proteins

Expi293 cells (RRID:CVCL_D615, ThermoFisher) for protein expression were maintained at 37 °C and 8% CO_2_ in Expi293 expression medium (ThermoFisher). Transfections were performed with an Expi293 Expression System Kit (ThermoFisher) and maintained under shaking at 130 rpm. Expression vectors encoding RBDs from spike proteins in SARS-2 Beta (GenBank QUT64557.1), SARS-2 WA1 (GenBank MN985325.1), SARS-2 BA.5 (GenBank UPN16705.1), SARS-2 BA.2.86 (EPI_ISL_18125249), SARS-2 XBB.1.5 (GenBank UZG29433.1), SARS-2 JN.1 (WPF38074.1), SARS-2 KP.3,^45^ RaTG13-CoV (GenBank QHR63300), SHC014-CoV (GenBank KC881005), Rs4081-CoV (GenBank KY417143), pangolin17-CoV (GenBank QIA48632), RmYN02-CoV (GSAID EPI_ISL_412977), Rf1-CoV (GenBank DQ412042), WIV1-CoV (GenBank KF367457), Yun11-CoV (GenBank JX993988), BM-4831-CoV (GenBank NC014470), BtKY72-CoV (GenBank KY352407), Khosta-2 CoV (QVN46569.1), and SARS-1 (GenBank AAP13441.1) were constructed as described^14,15,46^ to include a C-terminal tag (6xHis; G-HHHHHH) and SpyTag003 (RGVPHIVMVDAYKRYK)^47^ (for coupling of SARS-2 Beta, RaTG13, SHC014, Rs4081, pangolin17-CoV, RmYN02, Rf1, and WIV1 RBDs to SpyCatcher003-mi3) or a 15-residue Avi-tag (GLNDIFEAQKIEWHE) followed by a 6xHis tag for ELISAs and systems serology profiling. RBDs were purified from transiently-transfected Expi293F cell (ThermoScientific) supernatants as described.^14^ Expression vectors encoding soluble sarbecovirus spike trimers with either 6P^48^ (SARS-2 WA1 and SARS-2 JN1) or 2P stabilizing mutations (SARS-1, Rf1, Rs4081, Yun11, Bm48-31) were constructed as described with a 15-residue Avi-tag (GLNDIFEAQKIEWHE) followed by a 6xHis tag for systems serology profiling.^48^ For systems serology, Avi-tagged RBDs and spikes were co-expressed with a plasmid encoding the BirA enzyme for in vivo biotinylation^49^ (kind gift of Michael Anaya, Caltech) using a 1:1 RBD to BirA plasmid ratio for transfections. Biotinylated RBDs and spike trimers were purified and stored as described.^13^

### Preparation of mosaic-8b RBD-nanoparticles

SpyCatcher003-mi3 nanoparticles^19^ were expressed in E. coli, purified, and incubated with a 2-fold molar excess (total RBD to mi3 subunit) of SpyTagged RBDs (equimolar mixture of eight RBDs) overnight at room temperature in Tris-buffered saline (TBS) as described.^14,15^ Conjugated RBD-mi3 particles and free RBDs were separated by size-exclusion chromatography (SEC) on a Superose 6 10/300 column (GE Healthcare) equilibrated with PBS (20 mM sodium phosphate pH 7.5, 150 mM NaCl). Mosaic-8b RBD-nanoparticle conjugations were evaluated by SDS-PAGE and Western blot as described.^14,15^ Concentrations of conjugated mosaic-8b RBD-nanoparticles are reported based on RBD content determined using a Bio-Rad Protein Assay. Mosaic-8b nanoparticles were aliquoted and flash frozen in liquid nitrogen before being stored at −80 °C.

### ALD preparation of mosaic-8b RBD-nanoparticles

Mosaic-8b RBD-nanoparticles were spray-dried in a buffer with trehalose and hydroxyethyl starch to form spherical microparticles ranging in diameter from ∼1 to 10 μm as measured by flow imaging microscopy.^4^ The microparticles were then added to a customized fluidized bed reactor to apply conformal alumina coats of the desired thickness. Each reactor cycle added one 2.3 Å molecular layer of alumina on the microparticle surface using sequential pulses of trimethyl aluminum vapor followed by water vapor in self-limiting reactions. The number of layers applied is linearly related to the antigen release (e.g., 50 layers has an in vivo release time of ∼1 week).

### Immunization of mice

Mouse procedures were approved by the Caltech Institutional Animal Care and Use Committee.

Female BALB/c mice (6 weeks-old) were purchased from Charles River Laboratories (RRID: MGI:2161072) and housed at Caltech for immunization studies. Animals were healthy upon receipt and were monitored before the study during a 7-day acclimation period. Mice were randomly assigned to experimental groups of 5-8 animals. Up to 5 mice were housed together. Cages were kept in a climate-controlled room (71-75 °F; 50 ± 20% relative humidity) and mice were fed PicoLab Rodent Diet 20 (LabDiet) ad libitum.

Prior to immunization, SD, ALD mix3, and ALD mix4 were each reconstituted in distilled water (for naïve mouse immunizations) or ultrapurified sesame oil (for pre-vaccinated mouse immunizations) and swirled continuously to ensure homogeneity. Mice in the naïve cohort were immunized intramuscularly (IM) at Day 0 with one dose of 5 µg mosaic-8b RBD-nanoparticles (RBD equivalent) of either SD, ALD mix3, or ALD mix4 in 100 µl (50 µL per hind leg) or with two bolus immunizations at Day 0 and Day 28 of conventionally delivered mosaic-8b RBD-nanoparticles as follows: 2.5 µg total of protein nanoparticle (RBD equivalents) per dose (5 µg total) in 100 µL (50 µL per hind leg) containing 50% v/v AddaVax adjuvant (InvivoGen).

For the pre-vaccinated cohort of mice, we used a Pfizer-like mRNA-LNP formulation for WA1 spike from Helix Biotech for the pre-vaccinations because we were unable to obtain licensed Pfizer-BioNTech or Moderna mRNA-LNP vaccines for research purposes from the respective companies. Mice were vaccinated IM with 1 µg of WA1 mRNA-LNP diluted to 50 µL in PBS at weeks −15 and −12. Mice were then immunized IM at Day 0 with one dose of 5 µg mosaic-8b RBD-nanoparticles (RBD equivalent) of either SD, ALD mix3, and ALD mix4 in 100 µL ultrapurified sesame oil (50 µL per hind leg), or two doses at Day 0 and Day 28 of conventionally delivered mosaic-8b RBD-nanoparticles as described above for the originally naïve cohort of mice (2 x 2.5 µg doses; 5 µg total).

### Serum ELISAs

Purified His-tagged RBD was diluted to 2.5 µg/mL in 0.1 M NaHCO_3_ pH 9.8, coated onto Nunc® MaxiSorp™ 384-well plates (Sigma), and incubated overnight at 4°C. Plates were blocked with 3% bovine serum albumin (BSA), 0.1% Tween 20 in TBS (TBS-T) for 1 hr at room temperature, after which the blocking solution was removed by aspiration. 1:100 dilutions of mouse serum were serially diluted by 3.1-fold in TBS-T/3% BSA and then added to the plates for 3 hours at room temperature, followed by washing with TBS-T. After a 1-hour incubation with a 1:100,000 dilution of HRP-conjugated anti-IgG secondary (goat anti-mouse IgG; Abcam; RRID: AB_955439), plates were washed with TBS-T, SuperSignal™ ELISA Femto Maximum Sensitivity Substrate (ThermoFisher) was added as per manufacturer’s instructions, and luminescence was read at 425 nm. Midpoint titers (ED_50_ values) were obtained using Graphpad Prism 10.1.0 to plot and analyze curves assuming a one-site binding model with a Hill coefficient.

### Pseudovirus neutralization assays

HEK293T cells (RRID:CVCL_0063) for pseudovirus production were cultured in Dulbecco’s modified Eagle’s medium (DMEM, Gibco) supplemented with 10% heat-inactivated fetal bovine serum (FBS, Bio-Techne), 1% Penicillin/Streptomycin (Gibco), and 1% L-Glutamine (Gibco) at 37 °C and 5% CO_2_. HEK293T-hACE2 cells (RRID:CVCL_A7UK)^50^ for neutralization assays were cultured in DMEM (Gibco) supplemented with 10% heat-inactivated FBS (Bio-Techne), 5 mg/mL gentamicin (Sigma-Aldrich), and 5 mg/mL blasticidin (Gibco) at 37 °C and 5% CO_2_.

Lentiviral-based viruses were prepared as described^51,52^ using genes encoding S protein sequences lacking C-terminal residues in the cytoplasmic tail: 21 residue (SARS-2 variants) or 19 residue cytoplasmic tail deletions (SARS-1, Khosta-2-SARS-1 chimera, BtKY72-SARS-1 chimera). BtKY72 (containing K493Y/T498W substitutions) and Khosta-2 pseudoviruses were made with chimeric spikes in which the RBD from SARS-1 (residues 323-501) was substituted with the RBD from BtKY72 K493Y/T498W (residues 327-503) or Khosta-2 (residues 324-500) as described.^53^ Cells were co-transfected with HIV-1-based lentiviral plasmids, a luciferase reporter gene, and a coronavirus spike construct, resulting in lentivirus-based pseudovirions expressing a sarbecovirus spike protein. Supernatants were harvested 48-72 hours post-transfection, filtered, and stored at - 80 °C. Pseudovirus infectivity was determined by titration using HEK293T-hACE2 cells.

For neutralization assays, pseudovirus was incubated with three-fold serially diluted sera from immunized mice for 1 hour at 37°C, then the serum/virus mixture was added to HEK293T-hACE2 target cells or high-hACE2 HEK-293T cell line expressing hACE2 encoded with a consensus Kozak sequence (for SHC014 assays; kindly provided by Kenneth Matreyek, Case Western Reserve University) and incubated for 48 hours at 37°C. After removing media, cells were lysed with Britelite Plus reagent (Revvity Health Sciences), and luciferase activity was measured as relative luminesce units (RLUs). Relative RLUs were normalized to RLUs from cells infected with pseudotyped virus in the absence of antiserum. Half-maximal inhibitory dilutions (ID_50_ values) were derived in AntibodyDatabase^54^ using 4-parameter nonlinear regression.

### Epitope mapping by DMS

DMS experiments to map epitopes recognized by serum Abs were performed in biological duplicates using independent mutant RBD yeast libraries (Beta,^55^ WA1,^56^ XBB.1.5,^57^ SARS-1,^58^ and PRD-0038^58^ generously provided by Tyler Starr, University of Utah) as described.^14,59^ In order to remove non-specific yeast-binding Abs, sera that had been heat inactivated for 30 min at 56 °C were incubated twice with 50 OD units of AWY101 yeast transformed with an empty vector. Libraries were induced for RBD expression in galactose-containing synthetic defined medium with casamino acids (6.7g/L Yeast Nitrogen Base, 5.0 g/L Casamino acids, 1.065 g/L MES acid, and 2% w/v galactose plus 0.1% w/v dextrose). After inducing for 18 hours, cells were washed twice and then incubated with serum for 1 hr at RT with gentle agitation. Cells were then washed twice and labeled for 1 hr with secondary Ab (1:200 Alexa Fluor-647-conjugated goat anti-mouse-IgG Fc-gamma, Jackson ImmunoResearch 115-605-008, RRID:AB_2338904).

Stained yeast cells were processed using a Sony SH800 cell sorter. Cells were gated to capture RBD mutants that had reduced Ab binding for a relatively high degree of RBD. For each sample, cells were collected until ∼5 × 10^6^ RBD^+^ cells were processed (corresponding to ∼5 × 10^5^-1 × 10^6^ RBD^+^ Ab-escaped cells. Ab-escaped cells were grown overnight in synthetic defined media (6.7 g/L Yeast Nitrogen Base, 5.0 g/L Casamino acids, 1.065 g/L MES acid, and 2% w/v dextrose + 100 U/mL penicillin + 100 µg/mL streptomycin) to expand cells prior to plasmid extraction. DNA extraction and Illumina sequencing were done as described.^60^ Raw sequencing data will be available on the NCBI SRA. Escape fractions were computed using previously described processing steps^59,60^ and implemented using a Swift DMS program available from authors upon request). Escape scores were calculated using a filter to remove variants with mutations that escaped binding because of poor expression, >1 amino acid mutation, or low sequencing counts as described^55,60^

Escape map visualizations shown as static line plots, logo plots, and structural depictions were created using Swift DMS as previously described.^60^ Line heights indicate the escape score for a particular amino acid substitution, calculated as described.^60^ In some visualizations, RBD sites were categorized based on epitope region,^22–25^ class 1 (pink) (RBD residues 403, 405, 406, 417, 420, 421, 453, 455-460, 473-478, 486, 487, 489, 503, 504); class 2 (purple) (residues 472, 479, 483-485, 490-495), class 3 (blue) (residues 341, 345, 346, 437-450, 496, 498-501,), class 4 (orange) (residues 365-390, 408), class 5 (orange) (residues 352-357, 396, 462-468). For structural depictions of DMS data, an RBD surface (PDB 6M0J) was colored by the site-wise escape metric at each site, with red scaled to be the maximum used to scale the y-axis. Residues exhibiting the highest escape fractions were highlighted with their residue number and colored according to epitope class.

We stratified DMS escape fraction values into four groups. Escape fractions for each RBD substitution range from 0 (no cells with this substitution were sorted into the escape bin) to 1 (all cells with this substitution were sorted into the serum Ab escape bin).^61^ The sum of escape fractions for all substitutions at a specific site is represented by the total escape peak.^61^ DMS profiles with total escape peaks <0.5 at all sites were classified as polyclass responses. DMS profiles with total escape peaks of 0.5 to 1, >1 to 2, or >2 at one or more sites were classified as weak, moderate, or strong escape profiles, respectively, corresponding to their RBD epitope (Table S1).

### Systems serology: Ab subclass and FcγR binding profiling

Serum samples from immunized mice were analyzed using modifications of a Luminex assay to quantify the levels of antigen-specific Ab subclasses and FcγR binding profiles.^62^ Briefly, avidin (Sigma-Aldrich Catalog #: A9275-25MG) was coupled to magnetic Luminex microspheres (LuminexCorp) by carbodiimide-NHS ester coupling, Sulfo-NHS (ThermoFisher™ Catalog number 24510) and 1-Ethyl-3-[3-dimethylaminopropyl]carbodiimide hydrochloride (EDC) (ThermoFisher™ Catalog number 22980) according to the manufacturer’s instructions. Avidin-coupled microspheres were blocked for 30 min with 1x assay buffer (1x PBS pH 7.4 diluted from 10x PBS (Invitrogen), 1% BSA) and then washed twice with the same buffer. Avidin-coupled microspheres were then loaded with a biotinylated antigen (a sarbecovirus spike or RBD) at room temperature for 2 hours in assay buffer and then blocked with 10 µM biotin (Millipore Sigma Catalog B4501-100MG). Antigen-loaded microspheres were then incubated with heat-inactivated serum samples for 1 h) at an appropriate sample dilution (1:250–1:1250 for IgG1, IgG2a, IgG2b, total IgG; 1:100 for IgG3, FcγR2b-binding IgGs, FcγR3-binding IgGs, FcγR4-binding IgGs) for 1 hour at room temperature in 96-well plates (Corning) with continuous shaking. Unbound Abs were removed by washing twice with assay buffer (200 µL per wash). Secondary Abs (Southern Biotech; PE-coupled anti-IgG1, IgG2a, IgG2b, IgG3) were added at a 1:1000 dilution in assay buffer and incubated for 1 h at room temperature with continuous shaking. Excess primary and secondary Abs were removed by washing twice with assay buffer (200 µL per wash). Beads were resuspended in 200 µL of assay buffer and run on a Luminex™ FLEXMAP 3D™ Instrument System. Median fluorescence intensity was calculated for all samples, which were run in duplicate.

For evaluation of the FcγR-binding IgGs, biotin-labeled soluble ectodomains of 6xHis-tagged FcγR2b, FcγR3, and FcγR4 were prepared by co-expression with BirA enzyme in Expi293T cells as described.^49^ Biotinylated FcγRs were purified on a HisTrap column (VWR) according to the manufacturer’s instructions and SEC and then bound to PE-streptavidin (eBioscience). Labeled FcγRs were then diluted in assay buffer (1:200) and incubated with serum-coated microspheres for 1 h at room temperature with continuous shaking. Unbound primary and PE-labeled FcγR were removed as described above. Beads were resuspended in 200 µL of assay buffer and run on a Luminex™ FLEXMAP 3D™ Instrument System. Median fluorescence intensity was calculated for all samples, which were run in duplicate.

Heatmaps of antigen-specific Ab responses (log₁₀-transformed) were generated using Python (v3.9.16) with the Pandas (v2.0.3), NumPy (v1.23.5), Matplotlib (v3.7.1), and Seaborn (v0.12.2) packages. Control proteins to evaluate non-specific binding (listed as “control” in Figure 6) were HIV-1 BG505 SOSIP Env^63^ for IgG1, IgG2a, IgG2b, and Total IgG, and soluble influenza A/California/04/09 hemagglutinin^64^ for IgG3, FcγR2b-, FcγR3-, and FcγR4-binding IgGs.

### Quantification and statistical analyses

Pairwise comparisons, a method to evaluate sets of mean binding titers against individual viral strains for different immunization cohorts, were used as described previously to determine whether results from different cohorts were significantly different from each other.^13^ Statistically significant titer differences between immunized groups for ELISAs were determined using analysis of variance (ANOVA) followed by Tukey’s multiple comparison post hoc tests with the Geisser-Greenhouse correction, with pairing by viral strain, of ED_50_s/ID_50_s (converted to log_10_ scale) calculated using GraphPad Prism 10.1.0. For neutralizing titers (Figure 2C, Figure 3C, Figure S1), statistically significant titer differences between immunized groups for each given strain were determined using ordinary one-way ANOVA followed by Tukey’s multiple comparison test, with single pooled variance.

For statistical analysis of systems serology results in Figure 6, responses were aggregated by computing the geometric mean across replicate samples for each immunogen-antigen combination. Log₁₀-transformed geometric means were compared across immunogen groups using Tukey’s multiple comparison post hoc tests with the Geisser-Greenhouse correction, with pairing by viral strain calculated using GraphPad Prism 10.1.0. Box-and-whisker plots were generated for each IgG subclass or FcγR-binding IgG, displaying individual antigen-level geometric means per immunogen. Statistical significance was annotated using brackets and asterisk notation (p < 0.05 *, p < 0.01 **, p < 0.001 ***, p < 0.0001 ****).

### Materials, data, and code availability

Materials generated in this study are available upon request through a Materials Transfer Agreement. Code used for data processing and visualization of DMS and systems serology results is available upon request.

Main text and supplementary data will be publicly available. DMS raw sequencing data will be available under an NCBI SRA. Materials are available upon request to corresponding authors with a signed material transfer agreement, and other information required to analyze the data in this paper is available from the lead contacts upon request. This paper does not report original code. The work is licensed under a Creative Commons Attribution 4.0 International (CC BY 4.0) license, which permits unrestricted use, distribution, and reproduction in any medium, provided the original work is properly cited. To view a copy of this license, visit https://creativecommons.org/licenses/by/4.0/. This license does not apply to figures/photos/artwork or other content included in the article that is credited to a third party; obtain authorization from the rights holder before using such material.

## ACKNOWLEDGEMENTS

We thank Holly Coleman and Amber Rauch for antigen spray-drying, Jesse Bloom (Fred Hutchinson Cancer Research Center) and Tyler Starr (University of Utah) for RBD libraries, Ryan McNamara and Galit Alter (Ragon Institute) for instructions for Systems Serology experiments, Jost Vielmetter, Luisa Segovia, Alyssa Player, Annie Lam, and the Caltech Beckman Institute Protein Expression Center for protein production, Igor Antoshechkin and the Caltech Millard and Muriel Jacobs Genetics and Genomics Laboratory for Illumina sequencing, Chengcheng Fan for making the mosaic-8b RBD-nanoparticle model used in figures, Anthony West for Swift DMS support, and Kaito Nagashima for critical reading of the manuscript. These studies were funded by Wellcome Leap (P.J.B.), the National Institutes of Health P01-AI165075 (P.J.B.), Gates Foundation INV-034638 (P.J.B.) and INV-002149/INV-042180 (R.L.G. and T.W.R.), and the Merkin Institute for Translational Research (Caltech). This manuscript is the result of funding in whole or in part by the National Institutes of Health (NIH). It is subject to the NIH Public Access Policy. Through acceptance of this federal funding, NIH has been given a right to make this manuscript publicly available in PubMed Central upon the Official Date of Publication, as defined by NIH.

## AUTHOR CONTRIBUTIONS

Conceptualization: A.A.C., J.R.K., T.W.R., R.L.G., P.J.B.; Methodology: A.A.C., J.R.K., S.R., H.H.F., T.W.R., R.L.G., P.J.B.; Software, A.J.G.; Investigation: A.A.C., J.R.K., A.V.R., S.R., A.-C.P.F., L.M., H.G., P.N.P.G.; Resources: H.G., H.H.F., T.W.R., R.L.G.; Writing – original draft: A.A.C., J.R.K., P.J.B.; Writing – review and editing: A.A.C., J.R.K., S.R., T.W.R., R.L.G., P.J.B.; Visualization: A.A.C., J.R.K., A.V.R., R.L.G.; Supervision: A.A.C., J.R.K., S.R., T.W.R., R.L.G., P.J.B.; Funding: T.W.R., R.L.G., P.J.B.

## References

1 Hou, X., Zaks, T., Langer, R. & Dong, Y. (2021). Lipid nanoparticles for mRNA delivery. Nat Rev Mater 6, 1078–1094.

2 Rappuoli, R., Alter, G. & Pulendran, B. (2024). Transforming vaccinology. Cell 187, 5171–5194.

3 Kim, E. H., Teerdhala, S. V., Padilla, M. S., Joseph, R. A., Li, J. J., Haley, R. M. & Mitchell, M. J. (2024). Lipid nanoparticle-mediated RNA delivery for immune cell modulation. Eur J Immunol 54, e2451008.

4 Garcea, R. L., Meinerz, N. M., Dong, M., Funke, H., Ghazvini, S. & Randolph, T. W. (2020). Single-administration, thermostable human papillomavirus vaccines prepared with atomic layer deposition technology. NPJ Vaccines 5, 45.

5 Dong, M., Meinerz, N. M., Walker, K. D., Garcea, R. L. & Randolph, T. W. (2021). Thermostability of a trivalent, capsomere-based vaccine for human papillomavirus infection. Eur J Pharm Biopharm 168, 131–138.

6 Witeof, A. E., Meinerz, N. M., Walker, K. D., Funke, H. H., Garcea, R. L. & Randolph, T. W. (2023). A Single Dose, Thermostable, Trivalent Human Papillomavirus Vaccine Formulated Using Atomic Layer Deposition. J Pharm Sci 112, 2223–2229.

7 Witeof, A. E., McClary, W. D., Rea, L. T., Yang, Q., Davis, M. M., Funke, H. H., Catalano, C. E. & Randolph, T. W. (2022). Atomic-Layer Deposition Processes Applied to Phage lambda and a Phage-like Particle Platform Yield Thermostable, Single-Shot Vaccines. J Pharm Sci 111, 1354–1362.

8 Coleman, H. J., Yang, Q., Robert, A., Padgette, H., Funke, H. H., Catalano, C. E. & Randolph, T. W. (2024). Formulation of three tailed bacteriophages by spray-drying and atomic layer deposition for thermal stability and controlled release. J Pharm Sci 113, 3238–3245.

9 Weimer, A. W. (2019). Particle atomic layer deposition. J Nanopart Res 21, 9.

10 Hassett, K. J., Meinerz, N. M., Semmelmann, F., Cousins, M. C., Garcea, R. L. & Randolph, T. W. (2015). Development of a highly thermostable, adjuvanted human papillomavirus vaccine. Eur J Pharm Biopharm 94, 220–228.

11 Koolaparambil Mukesh, R., Yinda, C. K., Munster, V. J. & van Doremalen, N. (2024). Beyond COVID-19: the promise of next-generation coronavirus vaccines. npj Viruses 2.

12 Wang, E., Cohen, A. A., Caldera, L. F., Keeffe, J. R., Rorick, A. V., Adia, Y. M., Gnanapragasam, P. N. P., Bjorkman, P. J. & Chakraborty, A. K. (2025). Designed mosaic nanoparticles enhance cross-reactive immune responses in mice. Cell 188, 1036–1050 e1011.

13 Cohen, A. A., Keeffe, J. R., Schiepers, A., Dross, S. E., Greaney, A. J., Rorick, A. V., Gao, H., Gnanapragasam, P. N. P., Fan, C., West, A. P., Jr., Ramsingh, A. I., Erasmus, J. H., Pata, J. D., Muramatsu, H., Pardi, N., Lin, P. J. C., Baxter, S., Cruz, R., Quintanar-Audelo, M., Robb, E., Serrano-Amatriain, C., Magneschi, L., Fotheringham, I. G., Fuller, D. H., Victora, G. D. & Bjorkman, P. J. (2024). Mosaic sarbecovirus nanoparticles elicit cross-reactive responses in pre-vaccinated animals. Cell 187, 5554–5571 e5519.

14 Cohen, A. A., van Doremalen, N., Greaney, A. J., Andersen, H., Sharma, A., Starr, T. N., Keeffe, J. R., Fan, C., Schulz, J. E., Gnanapragasam, P. N. P., Kakutani, L. M., West, A. P., Jr., Saturday, G., Lee, Y. E., Gao, H., Jette, C. A., Lewis, M. G., Tan, T. K., Townsend, A. R., Bloom, J. D., Munster, V. J. & Bjorkman, P. J. (2022). Mosaic RBD nanoparticles protect against challenge by diverse sarbecoviruses in animal models. Science 377, eabq0839.

15 Cohen, A. A., Gnanapragasam, P. N. P., Lee, Y. E., Hoffman, P. R., Ou, S., Kakutani, L. M., Keeffe, J. R., Wu, H. J., Howarth, M., West, A. P., Barnes, C. O., Nussenzweig, M. C. & Bjorkman, P. J. (2021). Mosaic nanoparticles elicit cross-reactive immune responses to zoonotic coronaviruses in mice. Science 371, 735–741.

16 Starr, T. N., Greaney, A. J., Hilton, S. K., Ellis, D., Crawford, K. H. D., Dingens, A. S., Navarro, M. J., Bowen, J. E., Tortorici, M. A., Walls, A. C., King, N. P., Veesler, D. & Bloom, J. D. (2020). Deep Mutational Scanning of SARS-CoV-2 Receptor Binding Domain Reveals Constraints on Folding and ACE2 Binding. Cell 182, 1295–1310.e1220.

17 Brune, K. D., Leneghan, D. B., Brian, I. J., Ishizuka, A. S., Bachmann, M. F., Draper, S. J., Biswas, S. & Howarth, M. (2016). Plug-and-Display: decoration of Virus-Like Particles via isopeptide bonds for modular immunization. Scientific reports 6, 19234.

18 Zakeri, B., Fierer, J. O., Celik, E., Chittock, E. C., Schwarz-Linek, U., Moy, V. T. & Howarth, M. (2012). Peptide tag forming a rapid covalent bond to a protein, through engineering a bacterial adhesin. Proc Natl Acad Sci U S A 109, E690–697.

19 Bruun, T. U. J., Andersson, A. C., Draper, S. J. & Howarth, M. (2018). Engineering a Rugged Nanoscaffold To Enhance Plug-and-Display Vaccination. ACS Nano 12, 8855–8866.

20 Francis, T., Jr. (1960). On the Doctrine of Original Antigenic Sin. Proceedings of the American Philosophical Society 104, 572–578.

21 Cobey, S. & Hensley, S. E. (2017). Immune history and influenza virus susceptibility. Current opinion in virology 22, 105–111.

22 Barnes, C. O., Jette, C. A., Abernathy, M. E., Dam, K.-M. A., Esswein, S. R., Gristick, H. B., Malyutin, A. G., Sharaf, N. G., Huey-Tubman, K. E., Lee, Y. E., Robbiani, D. F., Nussenzweig, M. C., West, A. P. & Bjorkman, P. J. (2020). SARS-CoV-2 neutralizing antibody structures inform therapeutic strategies. Nature 588, 682–687.

23 Jette, C. A., Cohen, A. A., Gnanapragasam, P. N. P., Muecksch, F., Lee, Y. E., Huey-Tubman, K. E., Schmidt, F., Hatziioannou, T., Bieniasz, P. D., Nussenzweig, M. C., West, A. P., Keeffe, J. R., Bjorkman, P. J. & Barnes, C. O. (2021). Broad cross-reactivity across sarbecoviruses exhibited by a subset of COVID-19 donor-derived neutralizing antibodies. Cell reports 36, 109760.

24 Jensen, J. L., Sankhala, R. S., Dussupt, V., Bai, H., Hajduczki, A., Lal, K. G., Chang, W. C., Martinez, E. J., Peterson, C. E., Golub, E. S., Rees, P. A., Mendez-Rivera, L., Zemil, M., Kavusak, E., Mayer, S. V., Wieczorek, L., Kannan, S., Doranz, B. J., Davidson, E., Yang, E. S., Zhang, Y., Chen, M., Choe, M., Wang, L., Gromowski, G. D., Koup, R. A., Michael, N. L., Polonis, V. R., Rolland, M., Modjarrad, K., Krebs, S. J. & Joyce, M. G. (2023). Targeting the Spike Receptor Binding Domain Class V Cryptic Epitope by an Antibody with Pan-Sarbecovirus Activity. J Virol 97, e0159622.

25 Cui, L., Li, T., Lan, M., Zhou, M., Xue, W., Zhang, S., Wang, H., Hong, M., Zhang, Y., Yuan, L., Sun, H., Ye, J., Zheng, Q., Guan, Y., Gu, Y., Xia, N. & Li, S. (2024). A cryptic site in class 5 epitope of SARS-CoV-2 RBD maintains highly conservation across natural isolates. iScience 27, 110208.

26 Crescioli, S., Jatiani, S. & Moise, L. (2025). With great power, comes great responsibility: the importance of broadly measuring Fc-mediated effector function early in the antibody development process. MAbs 17, 2453515.

27 Bruhns, P. & Jonsson, F. (2015). Mouse and human FcR effector functions. Immunol Rev 268, 25–51.

28 Ackerman, M. E., Barouch, D. H. & Alter, G. (2017). Systems serology for evaluation of HIV vaccine trials. Immunol Rev 275, 262–270.

29 Hogenesch, H. (2012). Mechanism of immunopotentiation and safety of aluminum adjuvants. Front Immunol 3, 406.

30 Brubaker, S. W., Walters, I. R., Hite, E. M., Antunez, L. R., Palm, E. L., Funke, H. H. & Steadman, B. L. (2024). Demonstration of Tunable Control over a Delayed-Release Vaccine Using Atomic Layer Deposition. Vaccines (Basel) 12.

31 Li, C., Lee, A., Grigoryan, L., Arunachalam, P. S., Scott, M. K. D., Trisal, M., Wimmers, F., Sanyal, M., Weidenbacher, P. A., Feng, Y., Adamska, J. Z., Valore, E., Wang, Y., Verma, R., Reis, N., Dunham, D., O’Hara, R., Park, H., Luo, W., Gitlin, A. D., Kim, P., Khatri, P., Nadeau, K. C. & Pulendran, B. (2022). Mechanisms of innate and adaptive immunity to the Pfizer-BioNTech BNT162b2 vaccine. Nat Immunol 23, 543–555.

32 Gupta, A., Rudra, A., Reed, K., Langer, R. & Anderson, D. G. (2024). Advanced technologies for the development of infectious disease vaccines. Nat Rev Drug Discov 23, 914–938.

33 Nkolola, J. P. & Barouch, D. H. (2024). Prophylactic HIV-1 vaccine trials: past, present, and future. Lancet HIV 11, e117–e124.

34 Fan, C., Cohen, A. A., Park, M., Hung, A. F., Keeffe, J. R., Gnanapragasam, P. N. P., Lee, Y. E., Gao, H., Kakutani, L. M., Wu, Z., Kleanthous, H., Malecek, K. E., Williams, J. C. & Bjorkman, P. J. (2022). Neutralizing monoclonal antibodies elicited by mosaic RBD nanoparticles bind conserved sarbecovirus epitopes. Immunity 55, 2419–2435 e2410.

35 Fan, C., Keeffe, J. R., Malecek, K. E., Cohen, A. A., West, A. P., Jr., Baharani, V. A., Rorick, A. V., Gao, H., Gnanapragasam, P. N. P., Rho, S., Alvarez, J., Segovia, L. N., Hatziioannou, T., Bieniasz, P. D. & Bjorkman, P. J. (2025). Cross-reactive sarbecovirus antibodies induced by mosaic RBD nanoparticles. Proc Natl Acad Sci U S A 122, e2501637122.

36 Tamura, T., Irie, T., Deguchi, S., Yajima, H., Tsuda, M., Nasser, H., Mizuma, K., Plianchaisuk, A., Suzuki, S., Uriu, K., Begum, M. M., Shimizu, R., Jonathan, M., Suzuki, R., Kondo, T., Ito, H., Kamiyama, A., Yoshimatsu, K., Shofa, M., Hashimoto, R., Anraku, Y., Kimura, K. T., Kita, S., Sasaki, J., Sasaki-Tabata, K., Maenaka, K., Nao, N., Wang, L., Oda, Y., Genotype to Phenotype Japan, C., Ikeda, T., Saito, A., Matsuno, K., Ito, J., Tanaka, S., Sato, K., Hashiguchi, T., Takayama, K. & Fukuhara, T. (2024). Virological characteristics of the SARS-CoV-2 Omicron XBB.1.5 variant. Nat Commun 15, 1176.

37 Case, J. B., Sanapala, S., Dillen, C., Rhodes, V., Zmasek, C., Chicz, T. M., Switzer, C. E., Scheaffer, S. M., Georgiev, G., Jacob-Dolan, C., Hauser, B. M., Dos Anjos, D. C. C., Adams, L. J., Soudani, N., Liang, C. Y., Ying, B., McNamara, R. P., Scheuermann, R. H., Boon, A. C. M., Fremont, D. H., Whelan, S. P. J., Schmidt, A. G., Sette, A., Grifoni, A., Frieman, M. B. & Diamond, M. S. (2024). A trivalent mucosal vaccine encoding phylogenetically inferred ancestral RBD sequences confers pan-Sarbecovirus protection in mice. Cell Host Microbe 32, 2131–2147 e2138.

38 Tong, X., Wang, Q. X., Jung, W. Y., Chicz, T. M., Blanc, R., Parker, L. J., Barouch, D. H. & McNamara, R. P. (2024). Compartment-specific antibody correlates of protection to SARS-CoV-2 Omicron in macaques. Iscience 27.

39 Mackin, S. R., Desai, P., Whitener, B. M., Karl, C. E., Liu, M., Baric, R. S., Edwards, D. K., Chicz, T. M., McNamara, R. P., Alter, G. & Diamond, M. S. (2023). Fc-gammaR-dependent antibody effector functions are required for vaccine-mediated protection against antigen-shifted variants of SARS-CoV-2. Nat Microbiol 8, 569–580.

40 Irrgang, P., Gerling, J., Kocher, K., Lapuente, D., Steininger, P., Habenicht, K., Wytopil, M., Beileke, S., Schafer, S., Zhong, J., Ssebyatika, G., Krey, T., Falcone, V., Schulein, C., Peter, A. S., Nganou-Makamdop, K., Hengel, H., Held, J., Bogdan, C., Uberla, K., Schober, K., Winkler, T. H. & Tenbusch, M. (2023). Class switch toward noninflammatory, spike-specific IgG4 antibodies after repeated SARS-CoV-2 mRNA vaccination. Sci Immunol 8, eade2798.

41 Pillai, S. (2023). Is it bad, is it good, or is IgG4 just misunderstood? Sci Immunol 8, eadg7327.

42 Ben-Akiva, E., Chapman, A., Mao, T. & Irvine, D. J. (2025). Linking vaccine adjuvant mechanisms of action to function. Sci Immunol 10, eado5937.

43 Sievers, F., Wilm, A., Dineen, D., Gibson, T. J., Karplus, K., Li, W., Lopez, R., McWilliam, H., Remmert, M., Soding, J., Thompson, J. D. & Higgins, D. G. (2011). Fast, scalable generation of high-quality protein multiple sequence alignments using Clustal Omega. Mol Syst Biol 7, 539.

44 Landau, M., Mayrose, I., Rosenberg, Y., Glaser, F., Martz, E., Pupko, T. & Ben-Tal, N. (2005). ConSurf 2005: the projection of evolutionary conservation scores of residues on protein structures. Nucleic Acids Res 33, W299–302.

45 Kaku, Y., Yo, M. S., Tolentino, J. E., Uriu, K., Okumura, K., Genotype to Phenotype Japan, C., Ito, J. & Sato, K. (2024). Virological characteristics of the SARS-CoV-2 KP.3, LB.1, and KP.2.3 variants. Lancet Infect Dis 24, e482–e483.

46 Barnes, C. O., West, A. P., Jr., Huey-Tubman, K. E., Hoffmann, M. A. G., Sharaf, N. G., Hoffman, P. R., Koranda, N., Gristick, H. B., Gaebler, C., Muecksch, F., Lorenzi, J. C. C., Finkin, S., Hagglof, T., Hurley, A., Millard, K. G., Weisblum, Y., Schmidt, F., Hatziioannou, T., Bieniasz, P. D., Caskey, M., Robbiani, D. F., Nussenzweig, M. C. & Bjorkman, P. J. (2020). Structures of Human Antibodies Bound to SARS-CoV-2 Spike Reveal Common Epitopes and Recurrent Features of Antibodies. Cell 182, 828–842 e816.

47 Keeble, A. H., Turkki, P., Stokes, S., Khairil Anuar, I. N. A., Rahikainen, R., Hytönen, V. P. & Howarth, M. (2019). Approaching infinite affinity through engineering of peptide–protein interaction. Proceedings of the National Academy of Sciences 116, 26523–26533.

48 Hsieh, C. L., Goldsmith, J. A., Schaub, J. M., DiVenere, A. M., Kuo, H. C., Javanmardi, K., Le, K. C., Wrapp, D., Lee, A. G., Liu, Y., Chou, C. W., Byrne, P. O., Hjorth, C. K., Johnson, N. V., Ludes-Meyers, J., Nguyen, A. W., Park, J., Wang, N., Amengor, D., Lavinder, J. J., Ippolito, G. C., Maynard, J. A., Finkelstein, I. J. & McLellan, J. S. (2020). Structure-based design of prefusion-stabilized SARS-CoV-2 spikes. Science 369, 1501–1505.

49 Tykvart, J., Sacha, P., Barinka, C., Knedlik, T., Starkova, J., Lubkowski, J. & Konvalinka, J. (2012). Efficient and versatile one-step affinity purification of in vivo biotinylated proteins: expression, characterization and structure analysis of recombinant human glutamate carboxypeptidase II. Protein Expr Purif 82, 106–115.

50 Starr, T. N., Greaney, A. J., Addetia, A., Hannon, W. W., Choudhary, M. C., Dingens, A. S., Li, J. Z. & Bloom, J. D. (2021). Prospective mapping of viral mutations that escape antibodies used to treat COVID-19. Science 371, 850–854.

51 Crawford, K. H. D., Eguia, R., Dingens, A. S., Loes, A. N., Malone, K. D., Wolf, C. R., Chu, H. Y., Tortorici, M. A., Veesler, D., Murphy, M., Pettie, D., King, N. P., Balazs, A. B. & Bloom, J. D. (2020). Protocol and Reagents for Pseudotyping Lentiviral Particles with SARS-CoV-2 Spike Protein for Neutralization Assays. Viruses 12.

52 Robbiani, D. F., Gaebler, C., Muecksch, F., Lorenzi, J. C. C., Wang, Z., Cho, A., Agudelo, M., Barnes, C. O., Gazumyan, A., Finkin, S., Hagglof, T., Oliveira, T. Y., Viant, C., Hurley, A., Hoffmann, H. H., Millard, K. G., Kost, R. G., Cipolla, M., Gordon, K., Bianchini, F., Chen, S. T., Ramos, V., Patel, R., Dizon, J., Shimeliovich, I., Mendoza, P., Hartweger, H., Nogueira, L., Pack, M., Horowitz, J., Schmidt, F., Weisblum, Y., Michailidis, E., Ashbrook, A. W., Waltari, E., Pak, J. E., Huey-Tubman, K. E., Koranda, N., Hoffman, P. R., West, A. P., Jr., Rice, C. M., Hatziioannou, T., Bjorkman, P. J., Bieniasz, P. D., Caskey, M. & Nussenzweig, M. C. (2020). Convergent antibody responses to SARS-CoV-2 in convalescent individuals. Nature 584, 437–442.

53 Seifert, S. N., Bai, S., Fawcett, S., Norton, E. B., Zwezdaryk, K. J., Robinson, J., Gunn, B. & Letko, M. (2022). An ACE2-dependent Sarbecovirus in Russian bats is resistant to SARS-CoV-2 vaccines. PLoS Pathog 18, e1010828.

54 West, A. P., Jr., Scharf, L., Horwitz, J., Klein, F., Nussenzweig, M. C. & Bjorkman, P. J. (2013). Computational analysis of anti-HIV-1 antibody neutralization panel data to identify potential functional epitope residues. Proc Natl Acad Sci U S A 110, 10598–10603.

55 Greaney, A. J., Starr, T. N., Eguia, R. T., Loes, A. N., Khan, K., Karim, F., Cele, S., Bowen, J. E., Logue, J. K., Corti, D., Veesler, D., Chu, H. Y., Sigal, A. & Bloom, J. D. (2022). A SARS-CoV-2 variant elicits an antibody response with a shifted immunodominance hierarchy. PLoS Pathog 18, e1010248.

56 Starr, T. N., Greaney, A. J., Hannon, W. W., Loes, A. N., Hauser, K., Dillen, J. R., Ferri, E., Farrell, A. G., Dadonaite, B., McCallum, M., Matreyek, K. A., Corti, D., Veesler, D., Snell, G. & Bloom, J. D. (2022). Shifting mutational constraints in the SARS-CoV-2 receptor-binding domain during viral evolution. Science 377, 420–424.

57 Taylor, A. L. & Starr, T. N. (2023). Deep mutational scans of XBB.1.5 and BQ.1.1 reveal ongoing epistatic drift during SARS-CoV-2 evolution. PLoS Pathog 19, e1011901.

58 Lee, J., Zepeda, S. K., Park, Y. J., Taylor, A. L., Quispe, J., Stewart, C., Leaf, E. M., Treichel, C., Corti, D., King, N. P., Starr, T. N. & Veesler, D. (2023). Broad receptor tropism and immunogenicity of a clade 3 sarbecovirus. Cell Host Microbe 31, 1961–1973 e1911.

59 Greaney, A. J., Starr, T. N., Gilchuk, P., Zost, S. J., Binshtein, E., Loes, A. N., Hilton, S. K., Huddleston, J., Eguia, R., Crawford, K. H. D., Dingens, A. S., Nargi, R. S., Sutton, R. E., Suryadevara, N., Rothlauf, P. W., Liu, Z., Whelan, S. P. J., Carnahan, R. H., Crowe, J. E., Jr. & Bloom, J. D. (2021). Complete Mapping of Mutations to the SARS-CoV-2 Spike Receptor-Binding Domain that Escape Antibody Recognition. Cell Host Microbe 29, 44–57 e49.

60 Hills, R. A., Tan, T. K., Cohen, A. A., Keeffe, J. R., Keeble, A. H., Gnanapragasam, P. N. P., Storm, K. N., Rorick, A. V., West, A. P., Jr., Hill, M. L., Liu, S., Gilbert-Jaramillo, J., Afzal, M., Napier, A., Admans, G., James, W. S., Bjorkman, P. J., Townsend, A. R. & Howarth, M. R. (2024). Proactive vaccination using multiviral Quartet Nanocages to elicit broad anti-coronavirus responses. Nat Nanotechnol 19, 1216–1223.

61 Greaney, A. J., Starr, T. N., Barnes, C. O., Weisblum, Y., Schmidt, F., Caskey, M., Gaebler, C., Cho, A., Agudelo, M., Finkin, S., Wang, Z., Poston, D., Muecksch, F., Hatziioannou, T., Bieniasz, P. D., Robbiani, D. F., Nussenzweig, M. C., Bjorkman, P. J. & Bloom, J. D. (2021). Mapping mutations to the SARS-CoV-2 RBD that escape binding by different classes of antibodies. Nat Commun 12, 4196.

62 Brown, E. P., Dowell, K. G., Boesch, A. W., Normandin, E., Mahan, A. E., Chu, T., Barouch, D. H., Bailey-Kellogg, C., Alter, G. & Ackerman, M. E. (2017). Multiplexed Fc array for evaluation of antigen-specific antibody effector profiles. J Immunol Methods 443, 33–44.

63 Sanders, R. W., Derking, R., Cupo, A., Julien, J.-P., Yasmeen, A., de Val, N., Kim, H. J., Blattner, C., de la Peña, A. T., Korzun, J., Golabek, M., de los Reyes, K., Ketas, T. J., van Gils, M. J., King, C. R., Wilson, I. A., Ward, A. B., Klasse, P. J. & Moore, J. P. (2013). A Next-Generation Cleaved, Soluble HIV-1 Env Trimer, BG505 SOSIP.664 gp140, Expresses Multiple Epitopes for Broadly Neutralizing but Not Non-Neutralizing Antibodies. PLoS Pathogens 9, e1003618–1003620.

64 Cohen, A. A., Yang, Z., Gnanapragasam, P. N. P., Ou, S., Dam, K. A., Wang, H. & Bjorkman, P. J. (2021). Construction, characterization, and immunization of nanoparticles that display a diverse array of influenza HA trimers. PLoS One 16, e0247963.

